# Emergence of Heartbeat Frailty in Advanced Age: Perspectives from Life-Long EKG Recordings in Mice

**DOI:** 10.1101/2022.01.12.475659

**Authors:** Jack M Moen, Christopher H. Morrell, Ismayil Ahmet, Michael G. Matt, Moran Davoodi, Michael Petr, Shaquille Charles, Raphael deCabo, Yael Yaniv, Edward G Lakatta

## Abstract

SAN failure, aka sick-sinus syndrome, which features sinus bradycardia, SAN impulse pauses, and irregularity of RR interval rhythms are manifestations of SAN cell dysfunction that increases exponentially with advanced age, i.e., SAN frailty. Abnormalities in intrinsic RR interval variability may be the earliest signatures of SAN cell dysfunction leading to SAN frailty in late life. We measured RR interval variability within EKG timeseries prior to and during double autonomic blockade in long-lived C57/BL6 mice at 3 month intervals from 6 months of age until the end of life.

Long-lived mice (those that achieved the median cohort lifespan of 24 months and beyond) displayed relatively minor changes in intrinsic RR interval variability prior to 21 months of age. Between 21 and 30 months of age, marked changes in intrinsic RR interval variability signatures in time, frequency, non-linear, and fragmentation domains result in a marked increase in the mean intrinsic RR interval. The effects of autonomic input partially compensated for the prolongation of the mean RR interval by impacting the age-associated deterioration in the RR interval variability signatures toward a youthful pattern. Cross-sectional analyses of other subsets of mice at ages at or beyond the median life span of our longitudinal cohort demonstrated increased non-cardiac, constitutional, whole body frailty, a decrease in energetic efficiency, and an increase in respiratory exchange ratio. In this context, we interpret the progressive increase in intrinsic RR interval variability beyond 21 months of age to be an indication of heartbeat frailty.

## Introduction

The neuro-visceral autonomic axis continually conducts impulses at ms to sec time scales to and from the heart and other viscera via parasympathetic and sympathetic nerves. Beat to beat afferent signals (both neuronal and mechanical) originating within the heart signal to other parts of the heart and back out to the spinal cord, brain stem, and higher CNS structures, which in turn, elicit autonomic reflexes via efferent input of the central nervous system to the heart [1]. This autonomic input modulates the heart’s beating rate and rhythm by impacting the rate and rhythm of spontaneous action potential firing generated by mechanisms intrinsic to sinoatrial nodal (SAN) pacemaker cells. Although by convention, the number of heartbeats per minute is casually referred to as the *in vivo* heart rate (HR), analyses of successive R-R intervals within an EKG time-series detect substantial beat to beat interval variability, indicating that the sub-cellular and cell-wide mechanisms that underlie SAN impulse generation never achieve equilibrium [2, 3]

The combined influences of SAN pacemaker cell automaticity and its response to autonomic input determine the heart’s beating interval variability and mean beating rate. When the heartbeats in the absence of autonomic input, intrinsic base physiologic functions of SAN cells determine beat-to-beat interval variability from which the mean beating rate is calculated. SAN failure, aka “sick-sinus syndrome”, manifests as sinus bradycardia, SAN impulse pauses, and irregularity of rhythms of RR intervals within EKG time-series that result from SAN cell dysfunction and increases exponentially with advanced age often requiring electronic pacemaker implantation. Although the resting heart rate in humans changes very little beyond young adulthood, the mean **intrinsic** RR interval, measured during dual sympathetic and parasympathetic autonomic blockade, begins to decrease substantially between young adulthood and middle age [4], but has not been studied in humans of older ages. Because autonomic input to the SAN masks age-associated intrinsic dysfunction, intrinsic RR interval variability is likely to inform on the overall health status of the SAN in the absence of confounding effects from autonomic input, and changes in RR interval variability may be the earliest markers of SAN dysfunction, that ultimately progresses to SAN failure in advanced age. In short, **intrinsic** RR interval variability reflects the status of functions intrinsic to SAN cells that regulate the rate and rhythm of impulse emergence from the SAN.

The aging mouse manifests abnormalities in HR and heartbeat interval variability that recapitulate many of those described in humans [5–9]. **Cross-sectional** studies that compare average data in different mice that have survived to different ages indicate that the intrinsic heartbeat interval variability and mean intrinsic HR are relatively constant between young adulthood and middle age but become substantially altered later in life [7]. The increased sympathetic input to the SAN of older mice helps to reduce intrinsic heartbeat interval variability towards that of mice at younger ages. Other cross-sectional studies have associated changes in SAN automaticity in older mice with whole-body frailty [8, 10]. However, these cross-sectional studies lack **mouse-specific, longitudinal perspectives** on when and how changes in heartbeat interval variability leads to marked SAN dysfunction in individual long-lived mice, at an age when whole body frailty becomes manifest at older ages [11]. Longitudinal data on heart rhythm frailty in mice (or in humans) are mainly lacking because the longitudinal study design requires repeated measures of HRV that are implemented over a large part of the life course. The most important measures of cardiac frailty can be extended to mice, enabling the identification of SAN longevity markers in animals that live to or beyond the median life span.

We reasoned that changes in longitudinal measurements of intrinsic heartbeat interval variability in a given mouse will inform on the signature of late-life functional deterioration within and among SAN pacemaker cells. This functional deterioration leads to progressive changes in RR interval variability that increase the mean intrinsic RR interval in that mouse. To this end, we designed and implemented a longitudinal study that assessed heartbeat interval variability repeatedly in mice at 3-month intervals, beginning at 6 months of age and continuing to the end of life. We recorded EKG time-series at each age prior to (basal state) and during double autonomic blockade (intrinsic state) throughout the entire lifespan months in a cohort of 56 C57/BL6 black mice. This design permitted the elucidation of how the signature of mechanisms that underlie intrinsic SAN cell pacemaker function deteriorate over the life course. This design enabled the creation of the autonomic index, a novel set of measurements for the differences between mean basal and mean intrinsic state parameters measured at each age in a given mouse. This elucidated the rates at which autonomic signatures compensate for the deterioration in SAN intrinsic signatures during advanced age as the intrinsic SAN functions progress toward frailty. The unique nature of this experimental design enabled comparisons on the deterioration of intrinsic SAN function and compensatory autonomic signatures of heartbeat interval variability in advanced age to more traditional non-cardiac, whole body frailty indices, movement-based energetic efficiency, and energy substrate utilization.

## Methods

### Electrocardiogram

C57/BL6 mice, male, 3 months old (n=58), were obtained from Charles River Laboratories Inc. (Wilmington, MA). All studies were performed in accordance with *the Guide for the Care and Use of Laboratory Animals* published by the National Institutes of Health (NIH Publication no. 85-23, revised 1996). Experimental protocols were approved by the Animal Care and Use Committee of the National Institutes on Aging (protocol #441-LCS-2016).

Under light anesthesia with isoflurane (2% in oxygen) through nosecone at 0.4 ml/min, ECG electrode needles were inserted under the skin, a standard lead II ECG were recorded using Power Lab System (AD Instruments Inc.) at a sampling rate of 1,000 Hz. A heat lamp was positioned at a constant distance (25 cm) from the mouse during recordings to prevent heat loss. After let mice adapt to these conditions for twenty minutes, ECG was recorded continuously for 50 minutes: 10 minutes prior to, and 40 minutes after intraperitoneal injection of a saline solution (400 uL/30g body weight) with atropine (0.5 mg/kg) and propranolol (1.0 mg/kg). A representative timeseries of ECG RR interval recordings is illustrated in Supplemental Figure 1. ECG time-series were recorded in each mouse at three-month intervals, starting at six months of age, and continuing for the entire lifespan of each mouse.

The basal heart rate (HR) in each mouse was calculated as an average HR of 10 min recordings prior to drug and was reported as the mean basal RR interval (BRR). During the 40 min recordings after atropine and propranolol administration, HR was calculated as an average of each minute recordings, and the lowest HR was identified as the intrinsic HR (IHR). IHR is usually reached around 20 min after injection of atropine and propranolol. HR then returns to normal value around 40 min after injection. IHR reported as the mean intrinsic RR interval (IRR).

### EKG RR Interval Time-series Analyses

EKG time-series of RR intervals were analyzed using Physiozoo [12]. A mouse preset with rodent T waves was used to obtain a block average. Initially one-minute averages were obtained from the 10 minutes prior to injection and 40 minutes following injection. From the one-minute averages a block average was made for the set of basal recordings, and recordings in the presence of pharmacological autonomic blockade. Recordings of a single mouse were removed from the entire dataset, being at outlier with a resting heart rate >700 BPM.

Heart rate variability analysis also employed Physiozoo. ECGs were first analyzed to identify segments that fit strict selection criteria. This was mainly dependent on whether or not the heart rate was stationary for a sufficiently long period of time. Stationarity was determined based on the absence of linear trends, a high ectopic number, or a non-stable heart rate. Segments that met these eligibility criteria were included in the subset of tracings used for heart rate variability (for specific N numbers refer to Table S1). Generally, a 1.5-2-minute segment was selected, containing between 512 to 1,024 intervals. Rather, larger subsets were obtained for frequency domain analysis. The software was set up such that segments could be auto-analyzed based on length and segmented into similar sizes. Thus, our data was reported on files clipped to 512 intervals. After this we applied an automatic ectopic removal correction [12] in which any values outside of two standard deviations were removed.

### Statistics

All statistical analyses were implemented in R 3.2.3 [13] using RStudio [14].

### Mixed ANOVA Analyses

To test for age-associated differences in mean response, age was set as a factor under the assumption that recording intervals were approximately the same, and a mixed-effects ANOVA was conducted (Table S2).

### Linear mixed effects (LME) statistical analyses

A linear mixed effect (LME) model (lmerTest) was used to evaluate the effect of time following the initial measurement at 6 months of age on basal and intrinsic RR interval parameters, and autonomic index HR and HRV parameters.

### Estimating Mouse-Specific Rates of Change in RR Interval Variability

The average age trajectory is displayed using a loess smooth curve [15]. For many EKG time-series parameters, the trajectory over the age span is highly non-linear, exhibiting a sharp change around 21 months of age. This sharp change cannot be adequately modeled by a simple polynomial. Consequently, when needed, the trajectories are modeled in two parts: ages 6 – 21 months, and 21 to 30 months. To accommodate the curvature in the trajectories within each of these age-spans, a quadratic model in age is adopted, when needed. In this case, the full LME model becomes:

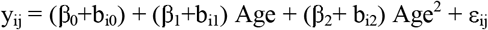

LME models contain both **fixed** and **random** terms. The β’s are the regression parameters for the **fixed-effects** variables, while b_i0_, b_i1_, and b_i2_ are the corresponding random effects. ε_i_ is the usual error term. The fixed-effects terms of LME models are the usual regression-type explanatory variables and their parameter estimates provide estimates of **average** rates of change in the response variable for changes in the associated variable. The **random effects** terms of LME models, however, allow for variability in initial value and trajectories among the mice. To obtain mouse-specific rates of change, age must now be considered as a numerical variable, and the trajectory over the age span must be appropriately modeled. The intercept random effect (b_i0_) allows for variability in individual mouse intercepts or starting points while the additional random effects, b_i1_ and b_i2_, allow for the linear and quadratic terms to vary among the animals.

In the full model, the rate of change in the response for individual mouse *i* is the derivative of the model function with respect to age and is given by:

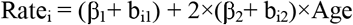

where the β’s and b_i_’s are replaced by the estimates obtained from the data. When terms are eliminated from the model, the appropriate remaining terms from the final model are used to obtain the mouse-specific rates of change. For many variables, the data could not accommodate the random effect for Age^2^. Consequently, in these cases, this random effect was removed from the model.

### Frailty Assessment

In order to compare the onset and severity of heartbeat frailty derived from our longitudinal EKG analyses, we conducted a cross-sectional study to assess traditional, non-cardiac specific, organism wide frailty in mice of advanced age.

### Frailty Index

We used a validated mouse frailty index consisting of non-cardiac factors [11]. Variability among mice in heart functions that change at older age have been shown to correlate better with this frailty index score than with chronologic age [16]. The frailty index was administered to a sub-set of older aged awake mice. A mouse Frailty Assessment Form was administered in parallel [6]. The assessment was performed in parallel by two different investigators in order to deter bias. The scores from each assessment were summed from the 31 parametric observations to calculate the mouse’s frailty score, and the scores determined by each investigator were averaged to generate final frailty score.

### Energetic Efficiency

Energetic efficiency, defined as kinetic work performed/oxygen consumption/unit time, was assessed cross sectionally in another sub-set of older mice. Work was measured as the distance traveled per unit of time and Energetic Efficiency. Oxygen consumption was measured through an enclosed treadmill coined as “metabolic treadmill”, where the mouse is subjected to run at its age groups’ average gait speed determined by over-ground walking analysis in earlier TSE MotoRater assessments. In order to ensure familiarity, mice were acclimated on the metabolic treadmills (Columbus Instruments International, Columbus, OH) at 5 m/min for 30 min the day prior to testing. The next day mice ran at the previously defined age-matched natural walking gait speed for 45 min [17]. The external motivators included electric shock from an electrified metal grid located near the moving belt to entice mice to run. Mice that were unwilling or incapable to continue running after being shocked 5 consecutive times within a few seconds met criterion for exhaustion and the testing ends.

## Results

### Longitudinal Cohort Longevity

The Kaplan-Meier Survival Curve of our entire cohort of C57/BL6 is shown in Fig 1A. The median life span of the total C57/BL6 cohort (n = 58) of our study was approximately 24 months (Fig 1). Because the main focus of our study was to identify how HR and HRV become altered in advanced age, we analyzed the life-long (6-30 month) EKG time series of each mouse in the basal and intrinsic states. We took 30 months of age as the maximum total life span of our mouse cohort because only 3 of 58 (5.2% of the total cohort) survived to this age. We focused on long-lived (LL) mice (n = 29), which we defined as those that achieved the median cohort lifespan of 24 months. Thus, the number of LL mice that survived from 6 to 24 months of age is constant at all ages and that beyond 24 months of age, the number of surviving mice becomes reduced as aging progresses, i.e., n = 17, and 3 at 27 and 30 months respectively (Table S1).

**Figure 1.**
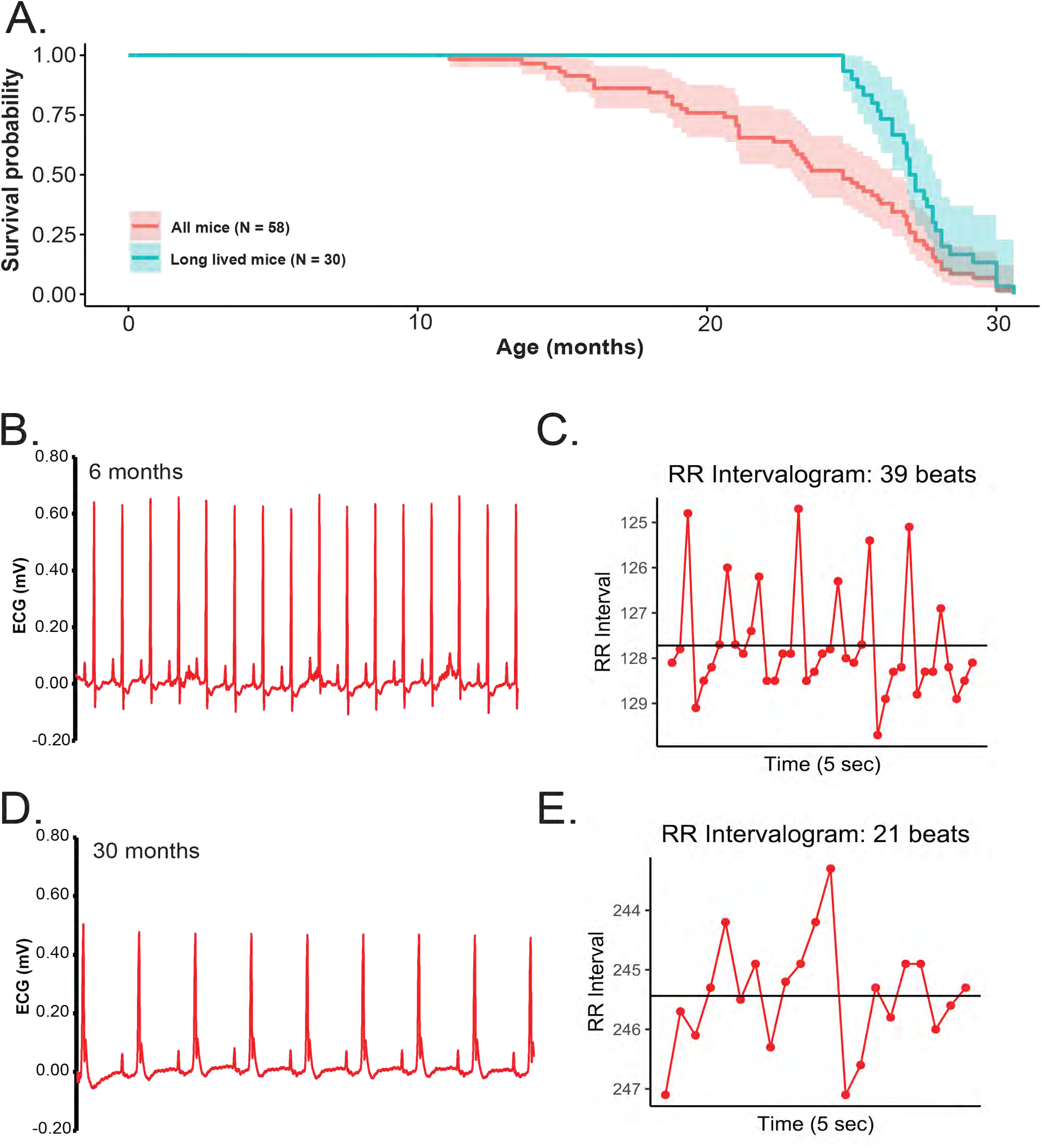
A. Kaplan-Meier Survival curves for entire cohort of mice (n = 58) and for long-lived subset of mice (n = 30). Intrinsic EKGs and Intervalograms at 6 months (B&C) and at 30 months (D&E).

The survival curve for the LL subset of our total mouse cohort is also shown in Fig 1A. Note its relative rectangularity compared to the survival curve of the entire cohort, and its steep rate of decline relative to that of the entire cohort The average life span of LL mice was 27.4 months (834 days or 119 weeks and 1 day) and relative to the entire cohort, LL mice at 24 months of age had already achieved about 75% of the mouse maximum life span.

### RR Interval Signature buried within EKG Time-Series

We assessed the EKG time-series RR interval variability patterns in the time, frequency, nonlinear, and fragmentation domains [18, 19]. Mean intrinsic RR interval variability data for all measured parameters at each age are listed in Table S1, and the corresponding mixed-effects ANOVA analyses of these data are presented in Table S2. The kinetics and degrees of synchronization within SAN cells [3] determine the intrinsic rhythms buried within an EKG time series during double autonomic blockade.

Representative EKG RR interval time-series recorded during autonomic blockade of a LL mouse at 6 and 30 months of age are shown in Figures 1B&D. RR interval variability (figures C&E) confirms that the SAN does not function as a metronome [20], as the heart beats in real time it has no a priori knowledge of its **mean** beating rate; instead, the time at which the next beat is generated is based on its memory of intrinsic mechanisms affected by prior beats [3]. Thus, a **mean** inter-beat within a time series does not exist in real-time but is calculated post hoc as the sum of individual and variable RR intervals in an EKG time series over a fixed period of time. For example, the mouse in Figure 1B at 6 months of age generated QRS complexes with 16 RR intervals, whereas at 30 months, only 8 RR intervals were generated within the same duration of time. The mean RR interval of an EKG time series does not capture this exquisite RR interval variability because the mean RR, calculated post hoc from RR intervals, assumes that all intervals are equal.

Non-constant interval variability among RR intervals within an EKG time series generates short and long-range correlations, nonlinearity components and variable frequency distributions of RR intervals within the time series [21]. The age-associated shifts in the RR interval variability pattern in the intrinsic state for the representative mouse (Figure 1B compared to Figure 1C, right panels) is striking, not only in the difference for the range of RR intervals but also differences in the synchronization of RR interval clusters. These RR interval rhythms buried within an EKG time series inform on the beat-to-beat variability in the kinetics of molecular mechanisms and the extent to which they are synchronized within and among cells [3].

### SAN Signatures within EKG Time Series Rhythms Measured Longitudinally Over the Life Course

The Poincaré diagram is a convenient, informative method to visualize frequency distributions of successive RR intervals within a time series that underlie the mean RR interval that relate to each other in the time domain. In a Poincaré diagram, a given RR interval (N) is plotted against the next RR interval (N+1). Fig 2A, inset, shows the Poincaré diagram of the LL mouse RR intervals presented in Fig 1. The points within the Poincaré plot can be fit to an ellipsoid, with the spread of the data measured as the SD1 or SD2;the Poincaré SD1 informs on short-range correlations among RR intervals within the EKG time-series, while SD2 informs on long-range correlations among intervals [22]. The intersection of SD1 and SD2 at the center of the ellipse is the mean RR interval, and the SD1: SD2 informs on the nonlinearity of the correlations of RR intervals within a given EKG time-series [22].

**Figure 2.**
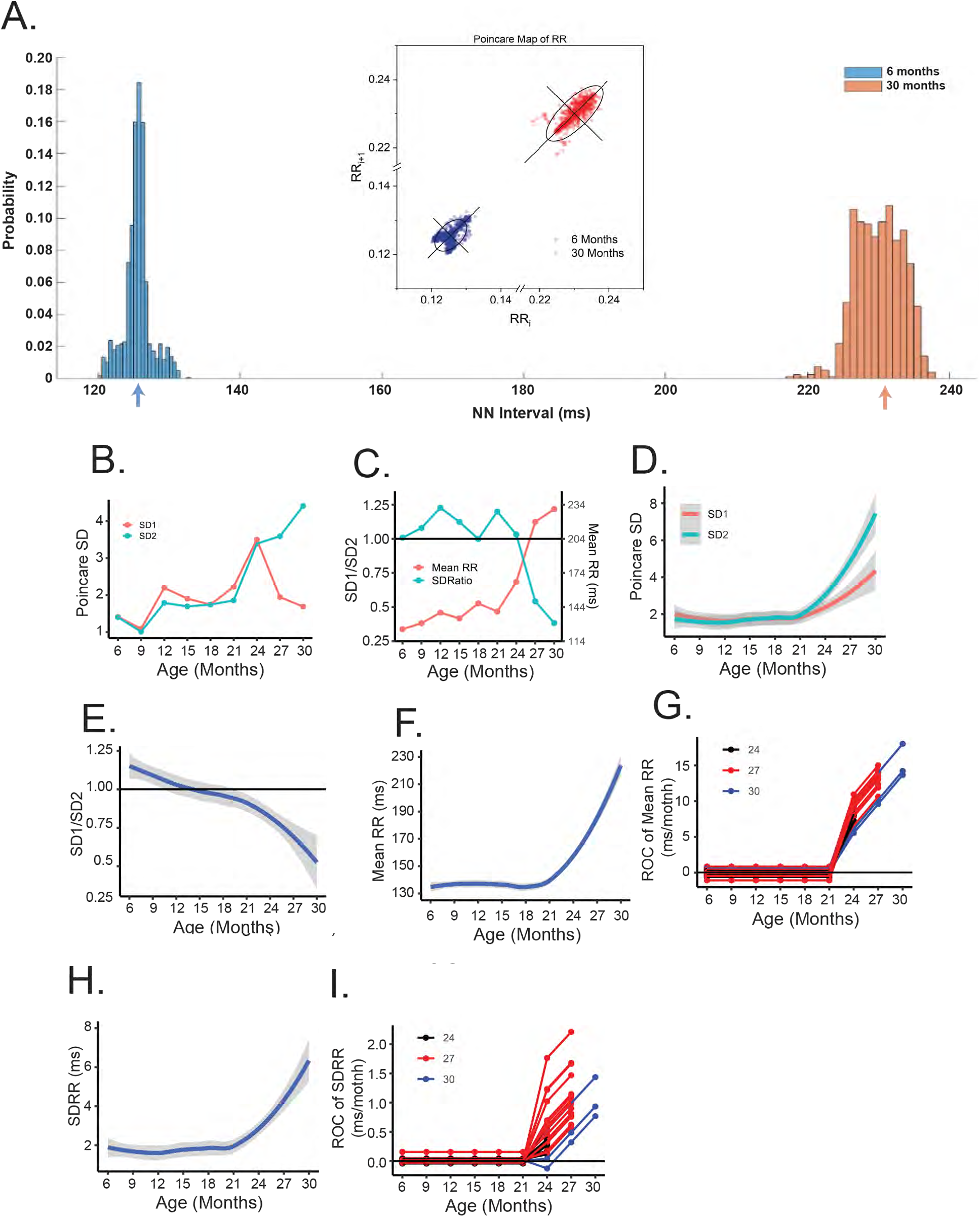
A. Synchronization within distributions of intrinsic EKG RR intervals at 6 and 30 months for mouse ZA1730. Inset, Poincare plot of intrinsic RR interval data at 6 and 30 months for mouse ZA1730. B. Intrinsic SD1 and SD2 for Mouse ZA1730. C. Intrinsic SD1/SD2 and Mean RR for Mouse ZA1730. D. Average loess smooth curves of intrinsic SD1 and SD2. E. Average loess smooth curves of intrinsic SD1/SD2. F. Average loess smooth curve for intrinsic mean RR. G. Mouse-specific rates of change for intrinsic mean RR in long-lived mice H. Average loess smooth curve of intrinsic SDRR. I. Mouse-specific rates of change of SDRR in long-lived mice.

Fig 2A shows the distributions of intrinsic RR intervals in the time domain (during autonomic blockade) for the same LL mouse at 6 and 30 months of age depicted in Fig 1 and Fig 2A inset. Note that not only is the RR interval (ms) distribution broader at 30 months than at 6 months of age, but the number of RR intervals within the distributions that are synchronized within narrow bins of intervals is substantially less at 30 months than at 6 months.

Intrinsic SD1, SD2, SD1:SD2, and the mean intrinsic RR interval throughout the life span of the LL mouse in Figs 1&2A are shown in Figs 2B&C. Note how the SD1 declines while SD2 increase beyond 21 months, creating a decreasing SD1:SD2. The average intrinsic SD1, SD2, and SD1:SD2 mean intrinsic RR intervals of all the LL cohort over its entire life span are shown in Figs 2D&E. On average, SD1, SD2 and the mean RR interval of LL mice markedly increased beyond 21 months of age, with the SD2 increasing to a much greater extent than SD1 (Fig 2D), reflecting nonlinearity (SD1:SD2) of short and long-range RR interval correlations as age increases beyond 21 months (Fig 2E). The intrinsic mean RR in all LL mice increases markedly beyond 21 months (Fig 2F).

Our experimental design permitted the calculation of mouse-specific rates of change (ROC) within aggregate RR interval data. In addition to providing the rate at which signatures change during a given mouse’s life course (i.e., mouse-specific rates), the ROC also provides insight into the rates at which inter-mouse-to-mouse variability of rates at which RR interval signatures change throughout the life course. ROC of intrinsic RR individual LL mice are illustrated in Fig 2G, the most significant increase occurring between 21 and 24 months of age.

SD of the mean RR interval (SDRR) can be calculated from the distribution of RR intervals. The **basal** SDRR has generally been attributed to the magnitude of parasympathetic input to the SAN [23], but the **intrinsic** SDRR measured during double autonomic blockade reflects beat-to-beat variability in mechanisms **intrinsic** to SAN cells. Beyond 21 months, the average increase in the intrinsic SDRR of the entire LL cohort (Fig 2H) is due to an increased age-associated beat-to-beat variability of kinetic functions that determine action potential firing rate within pacemaker cells of the SAN [3]. The ROC of intrinsic SDRR individual LL mice is illustrated in Fig 2I, the most significant increase occurring between 21 and 24 months.

Like the SD1:SD2, the slope coefficients α_1_ and α_2_ derived from detrended fluctuation analyses (DFA) together inform on nonlinearity of short-range DFA (slope coefficient α_1_) and long-range (DFA slope coefficient α_2_) RR interval correlations within an EKG time-series [21]. Examples of DFA analyses at 6 and 30 months in the same long-lived mouse depicted in Figures 1–2 are shown in Figure 3A. The calculated slope coefficients, α_1_ and α_2_, throughout the entire life course of this mouse are shown in Figure 3B, Tables S1 and S2. The average intrinsic α_1_ and α_2_ of the entire LL cohort also increased throughout the life span, with the increase accelerating sharply (nearly doubling) between 21 and 30 months (Fig 3C, Table S1, and S2). Mouse-specific ROC for α_1_ and α_2_ for all LL mice are shown in Figure 3D.

**Figure 3.**
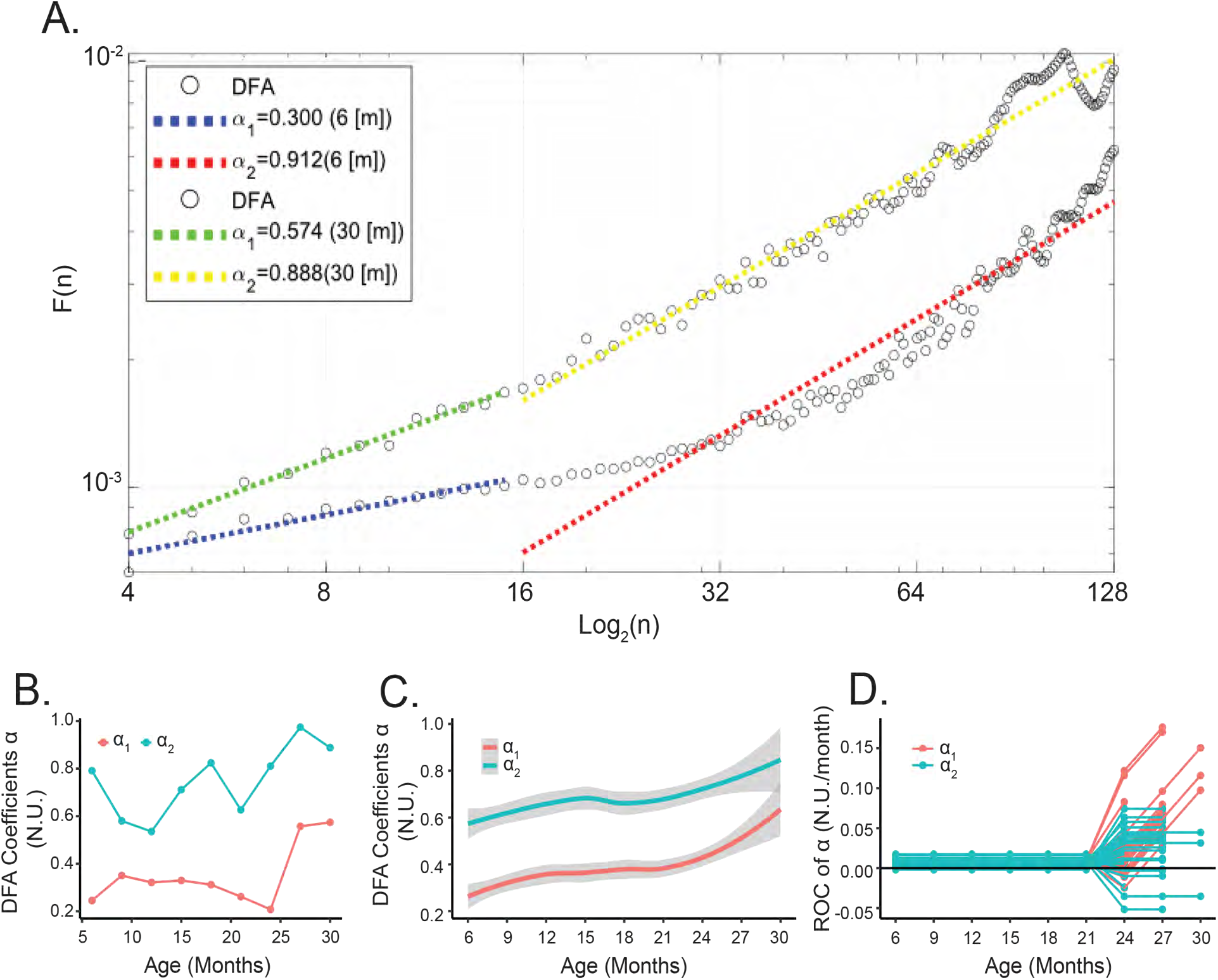
A. Detrended Fluctuation Analysis (DFA) illustrating intrinsic α_1_ and α_2_ at 6 and 30 months for mouse ZA1730. B. Trajectories of Intrinsic α_1_ and α_2_ in mouse ZA1730. C. Average loess smooth curves of intrinsic α_1_ and α_2_. D. Mouse-specific rates of change of intrinsic α_1_ and α_2_ in long-lived mice.

### SAN Aging Signature of RR Interval Rhythms in the Frequency Domain

Fast Fourier transforms provide power spectral densities (PSD) of RR intervals within an EKG interval time series, identifying different frequencies of hidden rhythms that are missed in EKG time-series analyses [24]. A greater total power informs on greater complexity, i.e., less coherence among RR intervals within the RR interval time series. Between 6 and 21 months, the mean intrinsic total power of the LL cohort did not vary significantly (Fig 4A, Table S2); but, beyond 21 months of age, the mean intrinsic total power **increased** by about 2-fold, similar to ROC for SDRR (Fig 2I). ROC of both total power and SDRR varied substantially among LL mice between 21 and 30 months (Fig 4B).

**Figure 4.**
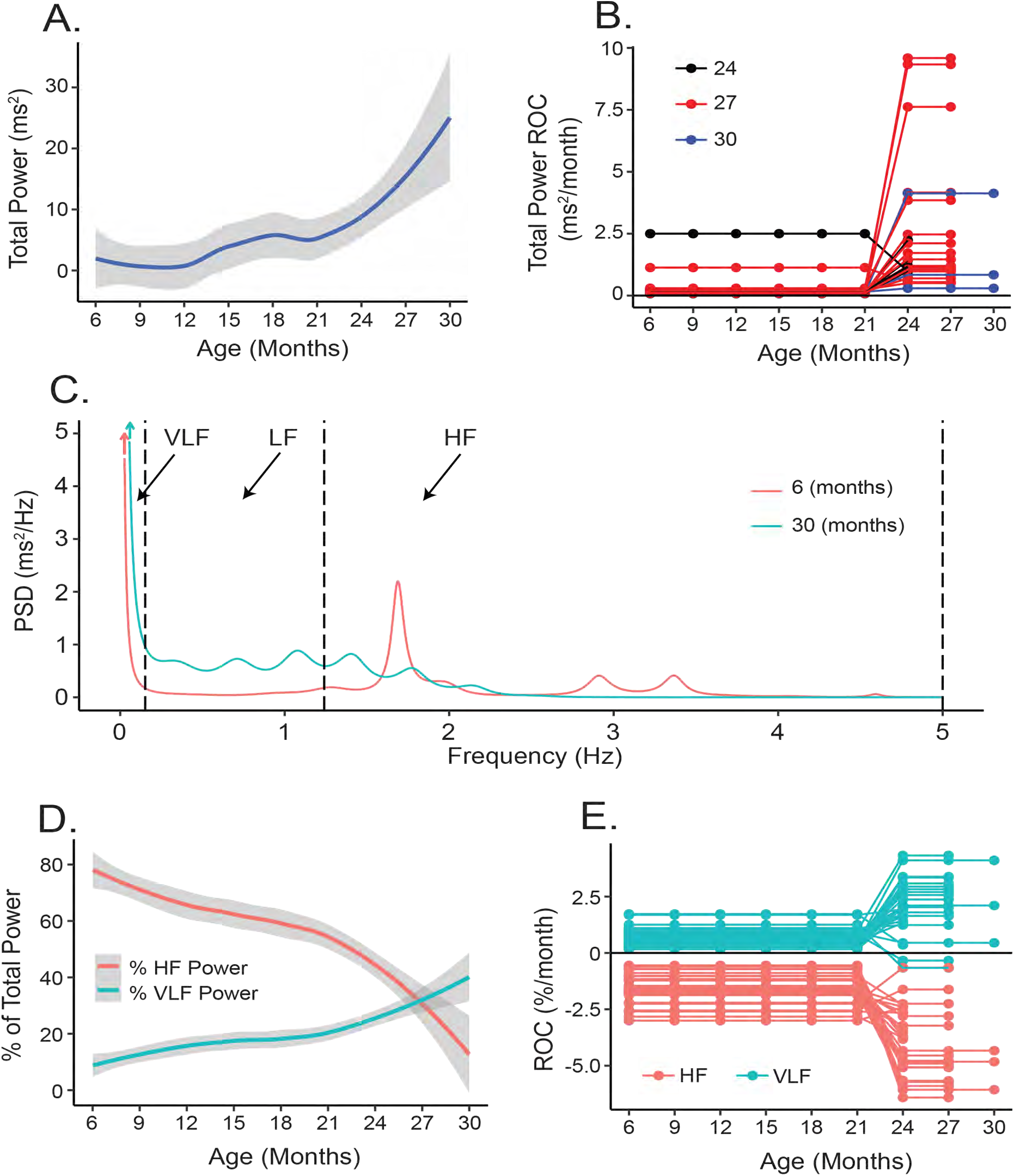
A. Average loess smooth curve of intrinsic Total Power (PSD). B. Mouse-specific rates of change of intrinsic Total Power (PSD) in long-lived mice. C. Intrinsic frequency domain data at 6 and 30 months for mouse ZA1730. D. Average loess smooth curves of % intrinsic VLF and HF. E. Mouse-specific rates of change of % intrinsic VLF and HF in long-lived mice.

The total power can be partitioned into regions that differ in their approximate frequencies, including the high frequency (HF), low frequency (LF), and very low frequency (VLF) power components (Fig 4C, Table S1). Examples of power spectra of RR intervals within the EKG time-series at 6 months and 30 months of age in the same long-lived mouse depicted in Figs 1–3, are shown in Fig 4C. Average values of intrinsic VLF power components normalized to total power in all mice increased modestly between 6 and 21 months (Fig 4D, Table S2). Beyond 21 months, the intrinsic %VLF power increased exponentially as intrinsic %HF power plummeted (Fig 4D, Table S2). Beyond 27 months, the **intrinsic** %VLF power exceeded that of intrinsic %HF power. The contribution of intrinsic HF power to intrinsic total power declined by about five-fold between 6 and 30 months in LL mice (Fig. 4D, Table S2).

### Intrinsic RR Interval Fragmentation Signatures

RR interval fragmentation analyses that attempt to quantify the erratic beating of the SAN by detecting small, aperiodic long-range fluctuations in RR intervals are an expansion of the aforementioned mentioned RR interval variability analyses [25]. RR interval fragmentation parameters assessed include: percentage of inflection points (PIP), or points at which the first RR difference changes sign; the percentage of short RR segments (PSS); the inverse of the average RR segment length (IALS); and the percentage of alternating RR segments (PAS). A larger value of each parameter denotes a more fragmented RR interval structure within the RR interval time series.

Changes in PIP, IALS, and PSS in our LL cohort resembled each other across the mouse lifespan (Fig S2). At ages less than 21 months, intrinsic IALS PIP and PSS displayed slight but statistically significant age-associated trends. Beyond 21 months, neither mean intrinsic PIP nor IALS varied significantly with age (Table S2); but, intrinsic PSS increased beyond 21 months (Fig S2, Table S2). The average LL cohort intrinsic PAS increased sharply beyond 21 months of age (Fig 5A). The individual ROC for the LL intrinsic PAS can be seen in Fig 5B. The Mean RR and PAS across the entire age span for the same mouse highlighted in Figs 1–4 are shown in Fig 5C.

**Figure 5.**
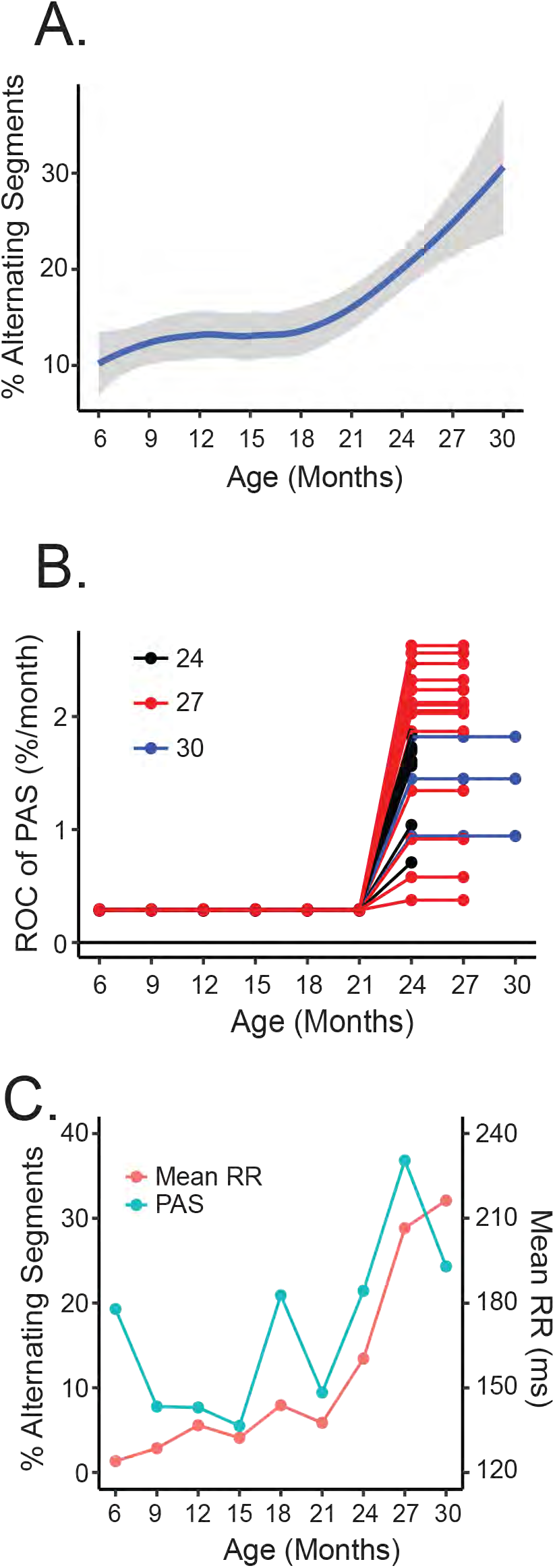
A. Average loess smooth curve of intrinsic PAS. B. Mouse-specific rates of change of intrinsic PAS in long-lived mice. C. Trajectories of Intrinsic PAS in mouse ZA1730.

### Relationships of the rates at which RR intervals change during the life course to the rates at which the mean RR interval changes

Because a crucial aspect of our RR interval variability analyses was to discover which intrinsic RR interval variability rhythms measured over the entire life course of a given mouse underlie the life course changes in the mean intrinsic RR interval in that mouse. In other terms, we sought to discover which RR interval variability signatures of aging in a given mouse define the rate at which the mean intrinsic RR interval increases with age in that mouse. We performed variable cluster analyses using mouse-specific ROC of all measured RR interval variability parameters between 21 and 30 months of age in each mouse. The dendrogram in Figure 6A illustrates the clustering of the mouse-specific ROC of the mean intrinsic RR interval and the intrinsic RR variability parameters between 21-30 months. ROC of intrinsic RR interval **variability** parameters that clustered with the ROC of mean intrinsic RR intervals in a given mouse (Fig 6A red cluster) were VLF, HF, α_2_, PAS, and SD1:SD2. Importantly, this cluster also includes bodyweight ROC, a component of the frailty assessment index (see below). Bivariate correlations for mouse-specific ROC of intrinsic RR variability signatures that clustered together throughout life (Fig 6A) are listed in Table 1 and illustrated in Fig S1. ROC of the intrinsic RR in individual mice as they age are illustrated in Fig 2G, with the most significant increase occurring between 21 and 24 months of age.

**Figure 6.**
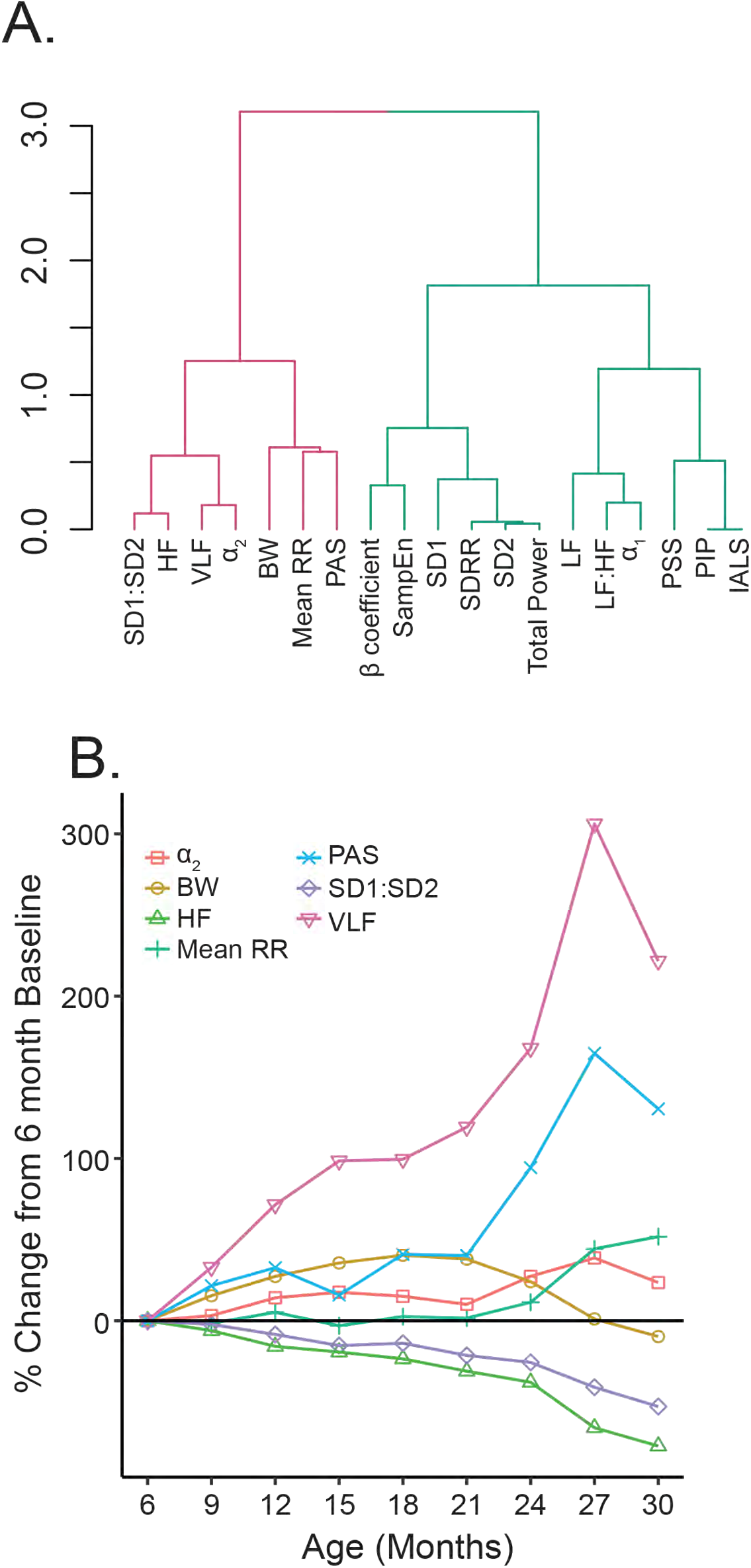
A. Cluster analysis dendogram of intrinsic rates of change in the late time period (after 21 months) in long lived mice. B. Relative change of intrinsic Mean RR and RR interval variability from 6 month baseline.

**Table 1.** Correlations and p-values among Intrinsic Rates of Change in long-lived mice

| | Body Weight | Mean RR | VLF | HF | SD1/SD2 | $\alpha_2$ | PAS |
| --- | --- | --- | --- | --- | --- | --- | --- |
| Body Weight: r |  | 0.0914 | 0.2669 | -0.1983 | -0.2627 | 0.3155 | 0.0779 |
| Body Weight: p |  | 0.6312 | 0.1539 | 0.2934 | 0.1608 | 0.0895 | 0.6823 |
| Mean RR : r | -0.1567 |  | -0.0091 | -0.0091 | 0.0601 | 0.1198 | 0.0603 |
| Mean RR : p | 0.4084 |  | 0.9621 | 0.9618 | 0.7524 | 0.5284 | 0.7515 |
| VLF : r | 0.2353 | 0.4774 |  | -0.8964 | -0.903 | 0.7772 | -0.3217 |
| VLF : p | 0.2106 | 0.0076 |  | 2.1E-11 | 8.7E-12 | 4.4E-07 | 0.0829 |
| HF : r | 0.2452 | -0.3914 | -0.7248 |  | 0.9507 | -0.5804 | 0.3271 |
| HF : p | 0.1915 | 0.0324 | 5.9E-06 |  | 8.9E-16 | 0.0008 | 0.0776 |
| SD1/SD2 : r | -0.1620 | -0.4264 | -0.7533 | 0.8818 |  | -0.6469 | 0.2817 |
| SD1/SD2 : p | 0.3924 | 0.0188 | 1.6E-06 | 1.2E-10 |  | 0.0001 | 0.1315 |
| $\alpha_2$ : r | -0.4078 | 0.4231 | 0.8189 | -0.4662 | -0.6444 | | -0.2243 |
| $\alpha_2$ : p | 0.0253 | 0.0198 | 3.2E-08 | 0.0094 | 0.0001 | | 0.2334 |
| PAS : r | -0.1567 | 0.4223 | 0.2074 | -0.2803 | -0.2927 | 0.1397 |  |
| PAS : p | 0.4084 | 0.0201 | 0.2716 | 0.1335 | 0.1165 | 0.4615 |  |

The upper triangle contains the correlations in the early time period (before 21 months) and the lower triangle contains the correlations in the late time period (after 21 months).

It is important to note that the ROC of RR interval variability signatures excluded from the red cluster (Fig 6A blue cluster) did **not** correlate with the ROC of the mean intrinsic RR interval between 21 and 30 months of age (Table S3). The age-associated increase in nonlinearity of RR interval variability (SD1:SD2) within short (SD1) and long (SD2) range RR interval correlations is a crucial factor of the marked increase in mean intrinsic RR interval prolongation between 21 and 30 months. Unlike the ROC SD1:SD2 (Table 1, Fig S3) neither the SD1, nor the SD2 ROC, per se, correlated with the age-associated increase in mean RR interval ROC (Table S3). Similarly, ROC for the standard deviation of the mean intrinsic RR intervals (SDRR) did not correlate with the ROC of the mean intrinsic RR during late life (Table S3). Unlike the SD1:SD2 (Table 1, S1), the SDRR does not provide information regarding the increase in the nonlinearity of RR interval variability within long and shortrange RR interval correlations that are **directly linked** to the rate at which the mean intrinsic RR interval increases in advanced age (Table 1, Fig S1).

The rate at which the mean intrinsic RR increases over time in a given mouse between 21 and 30 months was significantly correlated with concurrent increases in the rates at which α_2_ and %VLF power increased, as well as the rates at which %HF power and SD1:SD2 became reduced. The rates at which these RR interval parameters were underlying the increase in mean intrinsic RR interval during late-life were strongly correlated with each other: reductions in the rate of %HF power and SD1:SD2 are positively correlated, and the rates at which %VLF power and α_2_ increase are positively correlated (Table 1, Fig S1). Thus, ROC of relative %HF and %VLF power during the late-life course were inversely correlated with each other (Table 1).

Mouse-specific ROC of %HF and %VLF power were mirror images and therefore the rates at which they change are inversely related to each other. The rate at which the mean intrinsic RR increased over time in a given mouse between 21 and 30 months was significantly and positively correlated with concurrent increases in the rates at which α_2_ and %VLF power increased, and inversely with the rates at which %HF power and SD1:SD2 became reduced. The rates at which these RR interval parameters underlying the increase in mean intrinsic RR interval during late-life were strongly correlated with each other: reductions in the rate of %HF power and SD1:SD2 are positively correlated, as were the rates at which %VLF power and α_2_ increased (Fig S1, Table 1). Thus, ROC %HF and %VLF power were inversely correlated with each other during the late-life course (Table 1).

The ROC of intrinsic PAS (Fig 5B) in a given mouse was the only RR interval fragmentation index that correlated with the rate at which mean intrinsic RR interval increased during late life (Table 1, Fig S1).

Thus, between 21 and 30 months of age, the rate at which the mean intrinsic RR interval increased in a given mouse was associated with: (1) the rates at which HF and VLF components within the EKG time-series decrease and increase respectively; (2) positively with PAS; (3) increases in the rate at which nonlinearity of RR interval variability (SD1:SD2) between short-range (SD1) and long-range (SD2) RR interval correlations decreased; (4) the rates at which RR interval variability increased within long-range RR interval correlations (SD2, α_2_). Because the increase in α_2_ ROC in a given mouse was significantly correlated with the increase in mean intrinsic RR interval of that mouse (Tables 1 & S1), the ROC of α_2_ and SD1:SD2 during late life are inversely correlated with each other (Table 1, Fig S1).

### Quantitative Comparisons of the Relative Magnitudes of Life-Long Changes in RR Interval Variability Signatures and Mean RR Interval

To compare the relative magnitudes of changes in RR interval variability parameters (signature of aging in Table 1 and Fig 3A) that underlie the marked prolongation of the intrinsic RR in advanced age, we measured the values at 3-month intervals in each mouse, which were normalized to the 6 month values (Fig 6B). Intrinsic mean RR increased on average by 50% between 6 and 30 months, with much of the increase occurring beyond 21 months. Intrinsic %VLF power increased by about 2.5 fold; in contrast, intrinsic %HF power became reduced by about 20%, and the Poincaré SD1/SD2 was reduced by 50% between 6 and 30 months. PAS increased by 50% and DFA long-range intrinsic α_2_ increased by 25%. Note that beyond 21 months of age the increase in mean intrinsic RR interval is paralleled by sharp increases in α_2_, PAS, and %VLF power with the relative change in %VLF power greater than PAS, greater than α_2_; and by sharp reductions in SD1:SD2 and %HF power (beyond 24 months in Fig 6B).

In summation, our results show the impact of advanced age on the signature of RR interval variability and mean RR interval. The ROC of SD1: SD2, intrinsic HF, intrinsic VLF, and α_2_ in a given mouse are related to each other during late life, and these ROC of non-linear and frequency domain RR interval variability signatures underlie the mean intrinsic RR interval in advanced age.The unique clustering of mouse-specific RR interval variability signatures (Fig 6A) and correlations of these ROC with the rate at which the mean intrinsic RR interval in a given mouse increases in advanced age (Table 1, Fig S3) specifically inform on how the aging signatures of rhythms within EKG time series define the mean intrinsic RR interval as age advances.

### Impact of Autonomic Input on Age-Associated Changes in Intrinsic SAN RR Interval Signatures

Modulation of intrinsic heartbeat intervals imparted by autonomic input to the SAN reflects the combined effects of autonomic neurotransmitter impulses delivered to SAN cells and the responses of these pacemaker cells to that input. We next sought to investigate the effect of autonomic input (the difference between the mean intrinsic value and basal value of RR interval parameters in a given mouse) to glean additional insight into heartbeat dysfunction during late life. It is important to note that

Table S2 and Fig 7 show the signatures of autonomic input on RR interval variability signatures that underlie the rate at which the mean intrinsic of RR interval of LL mice increased during aging. Note that between 6 and 21 months of age, the difference between intrinsic and basal states of RR interval variability signatures did not differ (Tables 1 & S2). Between 21 and 30 months of age, the difference between intrinsic and basal state RR interval variability signatures differed markedly, during aging and among each other (Fig 7A-F, Table S2).

**Figure 7.**
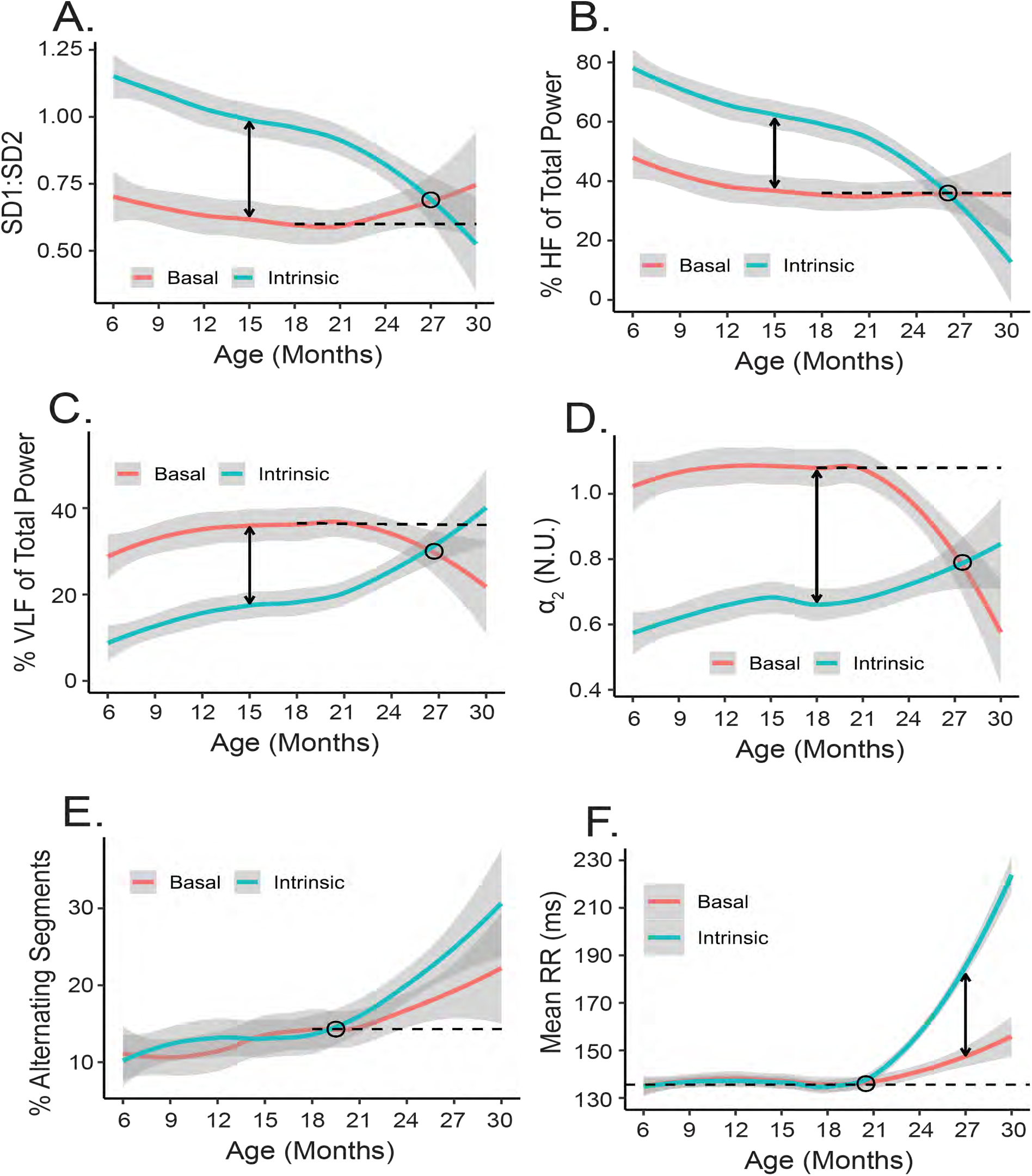
Average loess smooth curves for basal and intrinsic RR interval variablilty signatures and on the mean RR interval during the enitre life course of all long-lived mice for SD1:SD2 (A), %HF of total power (B), %VLF of total power (C), α_2_ (D), % alternating sgements (E), and mean RR (F). Arrows show the effect of autonomic modulation.

Before 21 months of age, SD1:SD2 was substantially less than the average SD1:SD2 of all mice in the intrinsic state (Fig 7A). Beyond 21 months, the average intrinsic SD1:SD2 began to decline while its basal state counterpart increased, with the average basal and average intrinsic SD1:SD2 converging at 27 months of age. Thus, up until 27 months of age, the effect of autonomic input was to reduce the intrinsic SD1:SD2. Note that average basal and intrinsic SD1 and SD2 in the three mice surviving up to 30 months of age crossed beyond 27 months, with the intrinsic SD1:SD2 reducing, while the basal SD1:SD2 increased, and that the crossing pattern of basal and intrinsic SD1:SD2 between 27 and 30 months of age is a general feature of other RR variability parameters that correlated with the mean RR intervals in very old mice e.g., %HF power (Fig 7B) %VLF power (Fig 7C) and DFA slope coefficient α_2_ (Fig 7D).

The effects that autonomic input imparts to %HF power (Fig 7B) is strikingly similar to that of its effect on the nonlinearity of short to long-range RR interval correlations within the time-series informed by SD1:SD2 (Fig 7A). Recall (Table 1, Fig S1) that the ROC of intrinsic %HF power and of SD1:SD2 in individual mice were positively correlated with each other. The effects of autonomic input on %VLF power (Fig 7C) and DFA α_2_ (Fig 7D) are also strikingly similar across the entire age span from 6 to 30 months. Recall also (Table 1, Fig S1) that: (1) the ROC for intrinsic %VLF power and α_2_ between 21 and 30 months were positively correlated to each other and to the rates at which the mean RR interval in individual mice changed over this age range; and (2) that the ROC for %VLF power and α_2_ were inversely correlated with the ROC of %HF power and SD1:SD2 (Table 1, Fig S2). Importantly, note in Figure 7 that the shapes of the mean %VLF power and α_2_ (Fig 7C-D) are also mirror images of those of %HF power and SD1:SD2 (Fig 7A-B); and also note the effects that autonomic signatures imparts to intrinsic %HF power and SD1:SD2 are strikingly similar.

PAS was the only RR interval fragmentation index throughout the entire life span that was not statistically significantly impacted by autonomic input (Table S2, S3). Average basal and intrinsic PAS in all LL mice beyond 21 months of age (Fig 7E) resembled a muted version of mean intrinsic and basal RR interval (Fig 7F), but unlike average intrinsic RR interval, the response to autonomic input was not significant. (Table S2). Although the average basal RR interval of all LL mice did not significantly differ between 6 and 21 months of age, the mean intrinsic RR manifested small but statistically significant changes (Table S2); and beyond 21 months of age, while average mean intrinsic RR precipitously increased (Fig 7F), autonomic input had a marked effect of reducing the mean RR interval (Table S2).

We may deduce that either autonomic input to the SAN increases, the responsiveness of SAN pacemaker cells to this input increases with age, or both mechanisms are involved. Because SAN responsiveness to autonomic neurotransmitters is known to be reduced in advanced age [26, 27], the reduction in mean intrinsic RR interval in late-life while autonomic input is intact reflects a **net** sympathetic response (increase sympathetic or reduced parasympathetic) to autonomic input, because of the directionality of the response (to reduce the RR interval or increase HR) (Fig 7F). Note, however, that beyond 21 months of age, autonomic input fails to restore the mean youthful basal RR interval that prevailed between 6 and 21 months (dashed line in Fig 7F), reflecting an inability of the net sympathetic autonomic input to overcome the high degree of disorder within the SAN tissue later in life. The impact of autonomic signatures on the intrinsic RR interval signatures over the entire life course of the representative mouse shown in Figs 1–7 are shown in Fig 8. Figs S2 & S3 illustrate the average basal and intrinsic RR intervals and RR variability parameters of the entire LL cohort having ROC that were not significantly correlated with mean RR interval ROC in individual mice.

**Figure 8.**
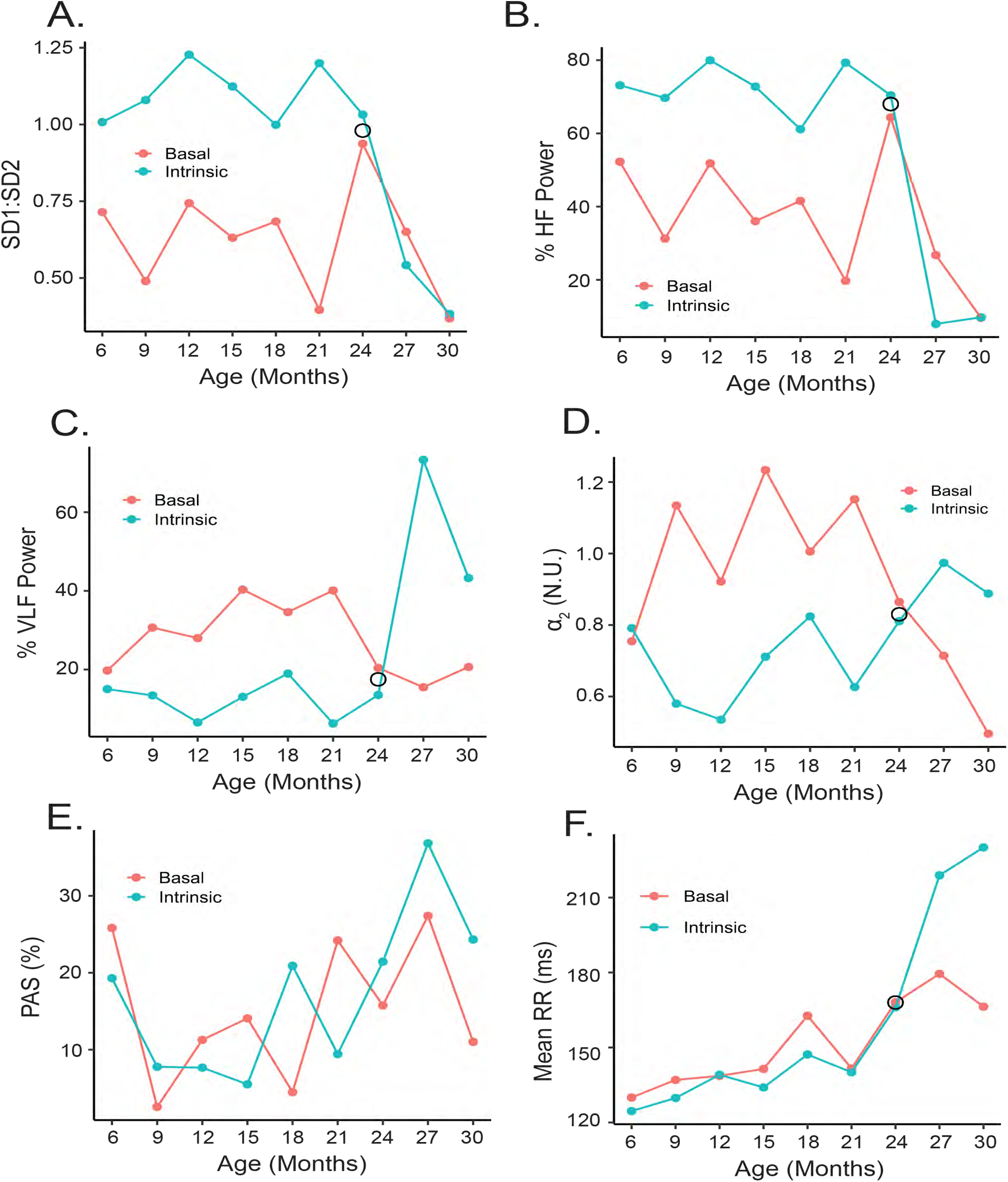
Impact of autonomic signature on intrinsic RR interval variability signatures; SD1:SD2 (A), %HF power (B), %VLF power (C), α_2_ (D), % PAS, and on the mean RR interval (F) during the entire lifespan of the representative mouse featured in Fig 1 – 6.

### Body Weight, Frailty Index and Energetic Efficiency

Loss of body weight is essential in non-cardiac frailty and may serve as an important biomarker in longevity [28]. Bodyweight in the long-lived mouse cohort changed in a non-linear manner, increasing significantly with age up to 18 months and significantly declining after 21 months until the end of life (Fig 9A), with the variability in mouse-specific ROC increasing beyond 18 months (Fig 9B). Considering body weight loss as a sign of frailty, it appears that the earliest signs of organismal frailty begin to emerge as early as 18 months (Fig 9A-B). Interestingly, the the ROC of the autonomic effect on intrinsic mean RR correlates (r = −0.3387) with the change in body weight.

**Figure 9.**
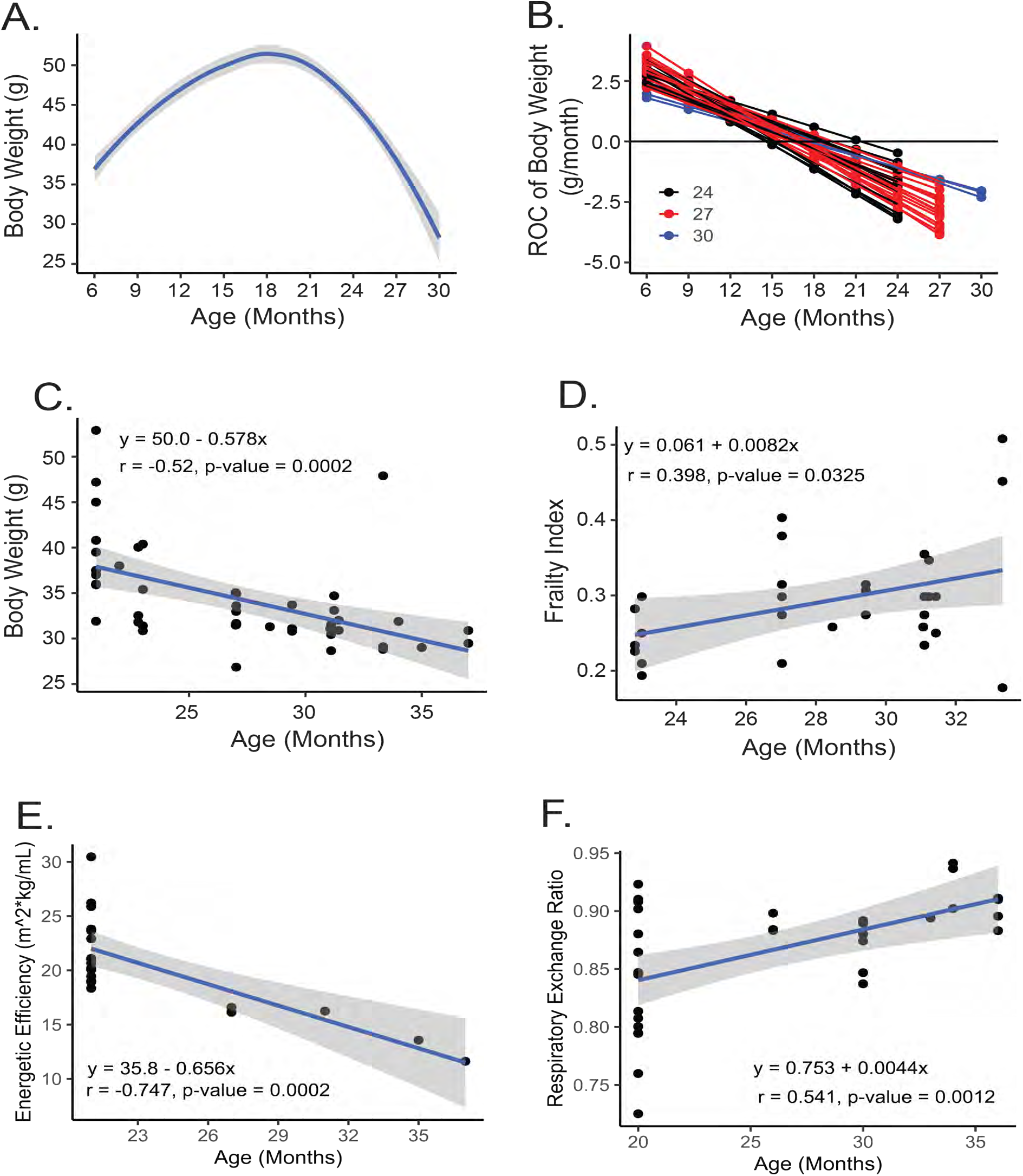
A. Average loess smooth curve for body weight. B. Mouse-specific rates of change of body weight. C. Body Weight. D. Frailty Index, E. Energetic efficiency, and F. Respiratory Exchange Ratio vs. Age.

We assessed body weight and whole organism frailty index that excluded cardiac parameters, in another cross-sectional sub-set of C57/BL6 mice at younger and older ages. Body weight increased substantially as age increased during late life (Fig 9C). As in our LL longitudinal cohort, bodyweight declined cross-sectionally in advanced age and was paralleled by an increase in frailty score (Fig 9D).

It is well known that the decline in health status is accompanied by a reduction in the efficiency of oxygen utilization [29, 30], which can be quantified as energetic efficiency measured as oxygen consumed during walking/speed of kinetic work performed [17]. Energetic efficiency measured cross-sectionally in another subset of mice also became reduced with advanced age (Fig 9E). An increase in the respiratory exchange ratio accompanied a reduction in energy efficiency (Fig 9F).

Thus, deterioration of the SAN interval variability signatures that underlie the marked increase in mean RR, i.e., heartbeat frailty observed during advanced age in our longitudinal cohort (Fig 1–8) are paralleled by increases in non-cardiac constitutional indices of frailty measured in other cross-sectional subsets (Fig 9). These additional perspectives on age-associated increases in the frailty index and reduced efficiency of O_2_ utilization establish that C57/BL6 mice at or beyond strain median life span of 24 months manifested constitutional signs of non-cardiac frailty.

## Discussion

We conducted a unique longitudinal study of mouse RR interval variability and mean RR interval analyses in EKG time-series recorded at 3-month intervals from 6 months to the end of life (approximately 30 months). By measuring RR interval variability prior to and during double autonomic blockade in a sub-set of mice that achieved the median cohort life span (24 months) or survived beyond this age we were able to assess how **intrinsic** SAN RR interval variability changed over the entire life course, particularly in advanced age. We specifically assessed mouse-specific rates of change for intrinsic RR interval variaility parameters to ascertain how these signatures vary within the same mouse over time, determining the rate at which the mean intrinsic RR interval increases during late-life in that mouse (SAN frailty), paralleling organismal frailty that emerges during late life.

### The Signatures of SAN Frailty in Long-Lived Mice of Advanced Age

Most measured RR interval variability parameters and mean RR interval remained stable up to about 18 months, but beyond 21 months of age changed dramatically. Between 21 and 30 months, most **intrinsic** RR variability parameters (measured during dual autonomic blockade) manifested progressive changes, reflecting dysfunction of mechanisms **intrinsic** to pacemaker cells residing within the SAN [3], causing the mean intrinsic RR interval to increase with advanced age.

Cluster analyses of mouse-specific rates of change (ROC) for the intrinsic RR interval variability correlations among longitudinal ROC of the clustered parameters enabled the identification of signature components of intrinsic RR interval variability that underlie the marked increase in the mean intrinsic RR interval that emerged beyond 21 months of age in LL mice (Table S2 Fig 6). These signatures of functional deterioration that occur within and among cells comprising SAN tissue underlying the marked increase in the mean intrinsic RR interval beyond 21 months of age are broadcast to the body surface as “heart beat music” [31] and can be heard on numerous EKG RR interval variability “channels,” including those that broadcast signals in the time, frequency, non-linear and fragmentation domains. Beyond 21 months of age cluster analyses pointed to ROC of intrinsic VLF, α2, HF, SD1/SD2 ratio and PAS being the signatures that sub-tended the rate at which the mean RR interval over this age range increased in individual mice.

The time at which successive RR intervals occur within a time series requires **memory** within SAN pacemaker cells created during prior beats [3]. Because each heartbeat is an emergent phenomenon [2], i.e., the SAN is not a metronome with a pre-written score. Increases of both intrinsic SD1 and SD2 means that RR interval variability within both short-range and long-range correlations of RR intervals generated within and among populations of SAN pacemaker cells become increased in advanced age in LL mice as SAN frailty begins to emerge. The nonlinearity of RR interval variability within short and long-range RR interval correlations also increase with aging, as reflcted in shifts in intrinsic SD1:SD2 over the late life course. A youthful profile of SD1, SD2, and SD1:SD2 likely requires high frequency signal processing, because the rate at which intrinsic %HF power is lost between 21 and 30 months strongly correlates with the rate at which intrinsic SD1:SD2 changes. Significant increases in SD1, SD2 and correlations between the rates at which intrinsic SD1:SD2 and %HF power components and decline in a given mouse are associated with the rate at which mean intrinsic RR interval increases in that mouse would appear to reflect, in part at least, a loss of memory with respect to when to generate the next AP as SAN frailty emerges. The corollary is that high frequency and nonlinearity of signals emerging from within and among populations of SAN cells informs on the memory that is required to maintain short (high frequency) and fairly regular (synchronized) RR intervals. The strong positive correlations of the rates at which HF components of RR intervals within the time-series decreased between 21 and 30 months in each mouse and the rates at which SD1:SD2 decreased in that mouse suggests that a reduction in the kinetics of pacemaker cell clock functions and desynchronization among these functions [3] may be linked to increased nonlinearity of RR interval variability within both short (SD1) and long-term (SD2) RR interval correlations. Rates at which intrinsic %HF power is lost and nonlinearity in (SD1: SD2) increase in a given mouse during advanced age were correlated with the rate at which mean intrinsic RR interval increased in that mouse.

The tight coupling between the rate at which the reduction in intrinsic %HF power and the rate at which intrinsic %VLF power increased, in the absence of a change in the rate of total power is likely attributable to a (Fig 7) reduction in the kinetics of intracellular Ca^2+^ and membrane potential transitions, leading to variability in mechanisms within and among SAN cells that relate to SAN pacemaker cell memory loss during the emergence of SAN frailty [3, 32, 33]. The shift in intrinsic HF power to VLF power of mice in advanced age is likely attributable to reductions in the kinetics of coupled-clock mechanisms that drive SAN pacemaker cell function [3, 33]. A reduction in these kinetics leads to desynchronization of molecular actions within SAN pacemaker cells associated with a loss of memory of synchronization effects created by the prior action potential [3].

The rates at which intrinsic DFA, slope coefficient α_2_, another measure of long term RR interval variability measure, and intrinsic %VLF power increased beyond 21 months of age in a given mouse were also significantly correlated with the rates at which the mean RR interval increases. Directionally opposite changes in the rate at which intrinsic α_2_ (increased) and SD1:SD2 decreased during advanced age means that the rate at which α_2_ increases reflects the rate at which HF signal processing is lost within and among populations of SAN cells. Thus, similar to the rate at which %VLF power increases beyond 21 months of age, the rate at which α_2_ increases is also correlated with the rate at which the mean intrinsic RR interval increases. The strong positive correlations between the rates at which %VLF power and α_2_ increased in a given mouse between 21 and 30 months of age indicate that the rate at which α_2_ increases reflects an increase in RR variability of VLF components within long-range RR interval (α_2_) correlations. In short, the most characteristic signature of SAN frailty is a loss in the ability of the SAN pacemaker cells to generate HF signals, resulting in reduced synchronization among the ensemble of cells. This translates into a markedly prolonged interval at which a given heartbeat follows the previous beat, resulting in a marked increase in the mean RR interval or reduction in mean intrinsic HR.

Increased SD1, SD2, SD1:SD2, increases in the DFA and slope coefficient α_2_, and shifts from intrinsic %HF power to intrinsic %VLF power underlie the increase in **intrinsic** percent alternating segments (PAS) within the EKG time-series of LL mice of advanced age that characterize SAN frailty beyond 21 months of age. This increase in the percentage of alternating long-short segments within the EKG interval time-series in LL mice of advanced age (i.e., increase in PAS) is reminiscent of action potential and calcium alternans that characterize dysfunctions within and among SAN pacemaker cells that underlie cardiac arrhythmias [34].

### Cellular and Molecular Basis of RR Intveral Variability Signatures of the Fraile SAN

Both experimental and theoretical perspectives have led to the idea that a coupled-oscillator system intrinsic to individual pacemaker cells drives normal, youthful automaticity [35]. Briefly, a chemical oscillator (“Calcium-Clock”) and an ensemble of current oscillators within the cell surface membrane (“Membrane Clock”) mutually entrain each other in a feed-forward manner to generate the rhythmic electrochemical gradients underlying action potential cycles [35]. Constitutive activation of adenylyl cyclase type 8 drives calcium and phosphorylation-dependent modulation of molecules that operate within the pacemaker cell clocks [36]. The effectiveness or fidelity of clock-coupling, determined by synchronization of mechanisms that regulate Ca^2+^ and membrane potential, controls the rhythm of action potential firing. The frail intrinsic RR interval variability signatures in LL mice of advanced age stem, at least in part, from disorder (desynchronization) among pacemaker cell clock molecules [3, 33], resulting in desynchronization of rhythms within and among clusters of pacemaker cells residing within SAN tissue [2].

Both age-associated reductions in clock molecule expression and reduced synchronization of molecular activation within individual pacemaker cells may be root causes of SAN failure in advanced age [33, 37]. But normal SAN function goes well beyond the function of individual pacemaker cells because rhythmic impulses that emerge from the SAN result from synchronization of heterogeneous, sub-cellular Ca^2+^ signals, not only within but also among SAN tissue resident pacemaker cells [2]. Thus, the initiation of each heartbeat within the SAN is an emergent phenomenon, and in addition to age-associated deficits within individual SAN cells, the ability to synchronize heterogeneous local signals among cells may be compromised in advanced age. Further, age-associated by pacemaker cell-matrix remodeling, including fibrosis [38] compounds the disordered state within the frail SAN.

### The Signatures of Autonomic Input to the Frail SAN

Adrenergic receptor activation increases cell calcium levels and protein phosphorylation, thereby increasing the entertainment of clock coupling [3]. When the effectiveness of clock-coupling increases in response to sympathetic autonomic input, RR variability and mean RR become reduced [36, 39]. Conversely, reduced effectiveness of clock coupling in response to parasympathetic autonomic input leads to prolonging the mean RR interval and increasing RR interval variability [3]. Thus, RR interval variability and the mean RR interval in the **basal** state result from modulation of the mechanisms intrinsic to SAN pacemaker cell that determine the **intrinsic** RR interval variability and mean RR interval.

Our results demonstrate that autonomic modulation of intrinsic RR interval variability differed over the life course, not only in magnitude but also in direction, particularly beginning around 21 months of age. As the rate of decline in intrinsic SD1:SD2 accelerated beyond 21 months of age, that of the basal SD1:SD2 increased with the basal and intrinsic values converging at 27 months of age. Thus, between 21 and 27 months, autonomic modulation of SAN pacemaker cells mechanisms markedly reduced the SD1:SD2 (Fig 7A), indicating that the impact of this autonomic input signature was to reduce the nonlinearity of RR interval variability that had developed within short to long-range intrinsic RR interval correlations in LL mice over this age range.

The signature of autonomic input that restored intrinsic %HF power was strikingly similar to the autonomic signature of SD1:SD2 between 21 and 27 months of age. Conversely, the signature of autonomic input was to reduce the increased intrinsic %VLF power and DFA α_2_, leading to preservation of their basal levels throughout the life course up to 27 months of age (Fig 7C-D). PAS was the only RR interval fragmentation parameter [40, 41] that did not have a statistically significant signature, i.e., was not significantly modified by autonomic input. The net result of these autonomic signatures on the intrinsic RR interval signatures was to reduce the basal RR interval to a more youthful level. Further, because the RR interval became reduced (i.e., HR was increased) this autonomic signature was net sympathetic in nature. Thus, age-associated changes in intrinsic RR interval signatures (heartbeat music broadcast to the body surface) reflected in the EKG time series is the net result of multiple changes within and among SAN cells and their responsiveness to autonomic input. Nevertheless, even after this autonomic input the basal state mean RR interval within a given mouse was not fully restored to its value at younger ages because the autonomic signatures could not fully restore the RR interval variability signatures to their youthful levels.

### The Relationship of SAN Frailty to Body-Wide Frailty

We showed in an additional cross-sectional C57/BL6 mouse cohort that over the age range in which marked disorder of SAN function emerged (in our LL longitudinal cohort beyond about 21 months of age), a bodywide frailty index not inclusive of cardiac parameters also sharply increased during late life (Fig 8D). An increased frailty score with advanced age has also been demonstrated in prior cross-sectional studies [17]. Of note, a prominent element within this frailty index is bodyweight loss, which not only occurred in our cross-sectional cohort, but also began to occur in all mice in our LL **longitudinal** cohort at 18 months of age (Fig 9). The increase in the frailty index and loss of body weight during late-life in our cross-sectional cohort was accompanied by a loss of energy efficiency (Fig 8E) and an increase in the respiratory exchange ratio (Fig 8F).

In summary, in the context of the presence of these other non-cardiac frailty markers in mice of advanced age, we may interpret the marked changes in increased intrinsic SAN RR variability that underlies the marked increase in the intrinsic mean RR interval that emerges in late life to be a signature of progressive SAN frailty. Specific aspects of this SAN frailty signature include: (1) increased non-linear intrinsic RR interval variability within short and long-range correlations of RR intervals; (2) reductions in the %HF power within intrinsic RR interval rhythms; (3) increase in %VLF intrinsic power within RR rhythms; (4) increased long-range detrended RR interval fluctuations; (5) progressive increase in the percentage of intrinsic RR interval segments exhibiting alternans. In toto, these RR interval variability signatures reflect an inability to generate high frequency RR intervals due to desynchronization of molecular functions within SAN pacemaker cells that require memory of prior RR intervals. The emergence and progression of this altered pattern of intrinsic RR interval variability beyond about 21 months of age cause the mean intrinsic RR interval to progressively increase over this age range. Autonomic modulation of this intrinsic SAN RR interval variability signature masks, to a large extent, SAN frailty cell that emerges during late life concurrently with the emergence of non-cardiac, body-wide, organismal frailty [42].

## Acknowledgements

This study was partially supported by the NIA Intramural Research Program and partially by ISF330/19. This research was partially supported by the Technion Hiroshi Fujiwara Cyber Security Research Center and the Israel Cyber Directorate (M.D).

We wish to acknowledge the indefatigable editorial assistance of Loretta E. Lakatta, R.N., B.S.N., and Syevda Tagirova Ph.D. for assistance with fitting Poincaré plot ellipses.

## Ethical Statement/Conflict of Interest

No author has any conflict of interest to report.

## Supplemental Figures

**Figure S1.**
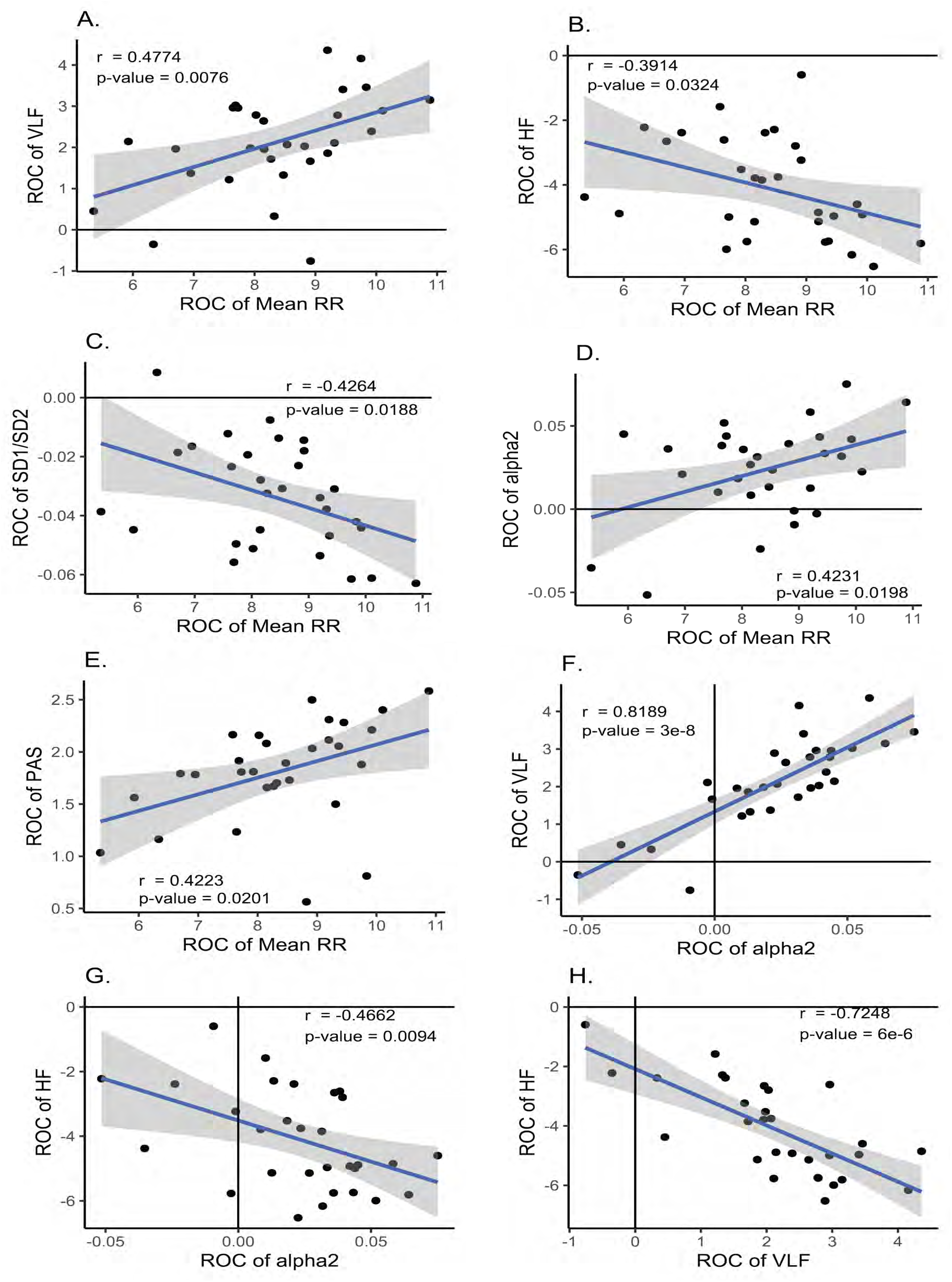

**Figure S2.**
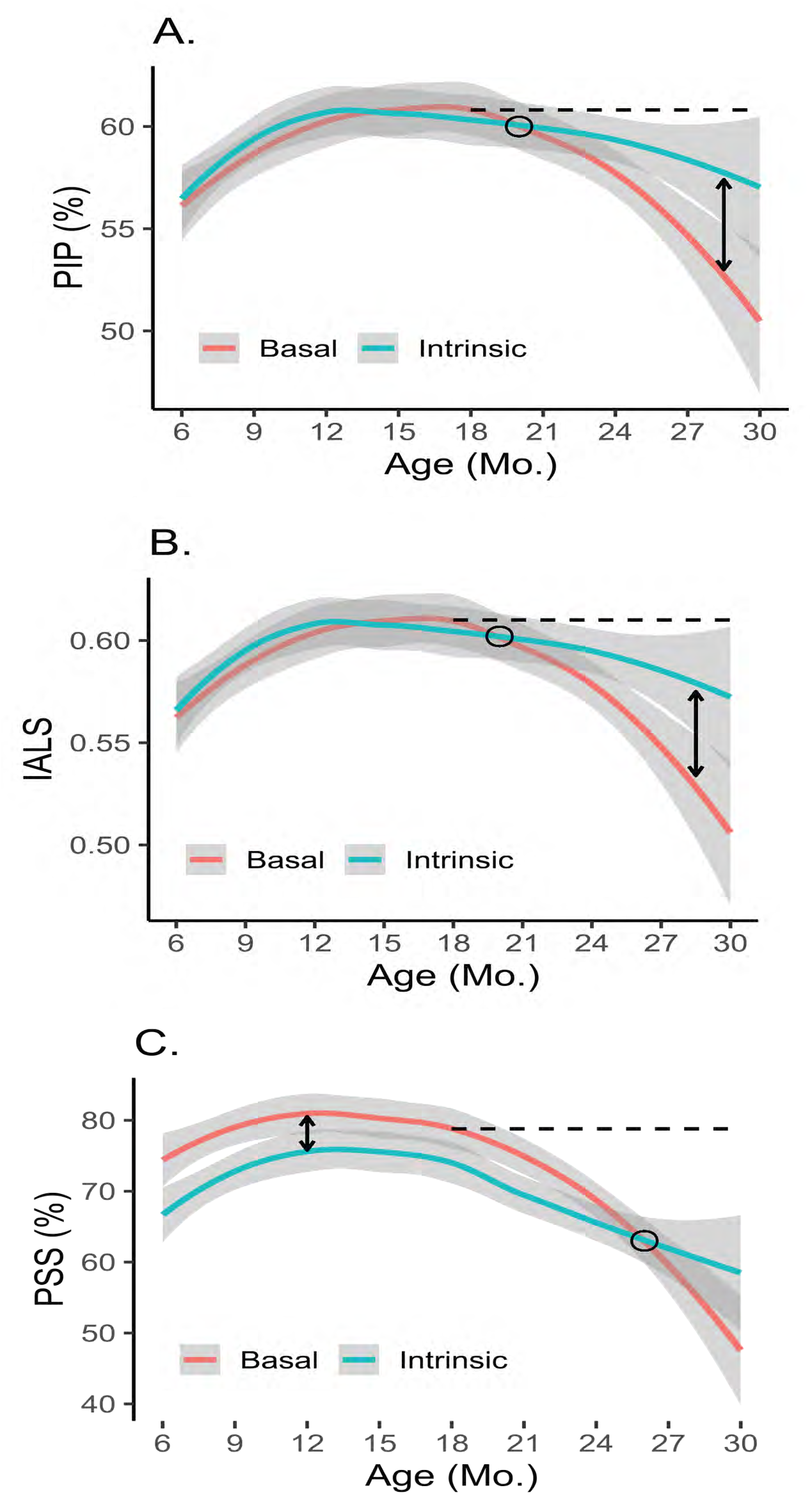

**Figure S3.**
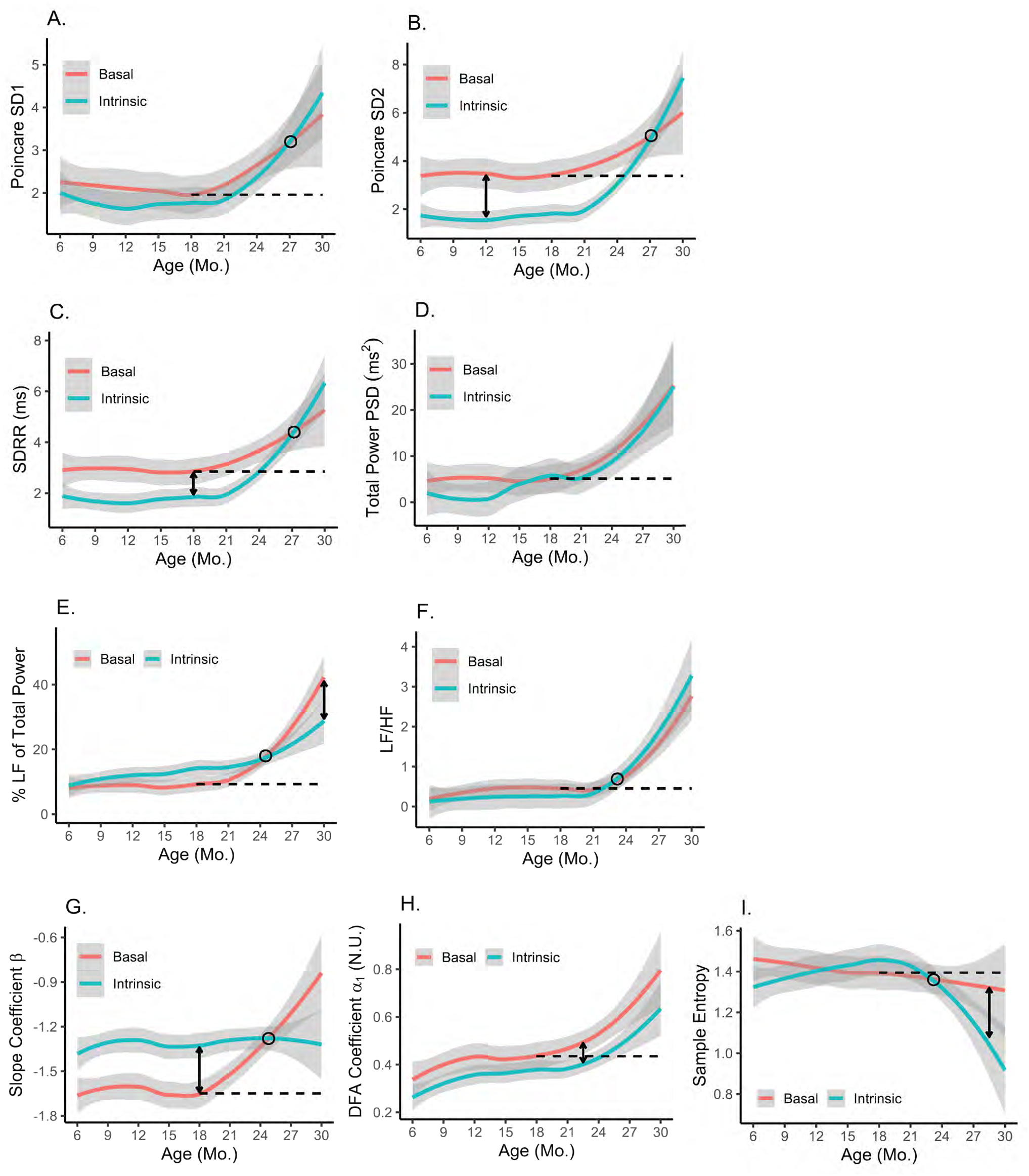

## Notes

### Competing Interest Statement

The authors have declared no competing interest.

### Summary of Updates

Two additional important citations have been included.

## References

1. Armour, J. A., Potential clinical relevance of the ‘little brain’ on the mammalian heart. Exp Physiol, 2008. 93(2): p. 165–76.

2. Bychkov, R., et al., Synchronized Cardiac Impulses Emerge From Heterogeneous Local Calcium Signals Within and Among Cells of Pacemaker Tissue. JACC Clin Electrophysiol, 2020. 6(8): p. 907–931.

3. Yang, D., et al., Ca2+ and Membrane Potential Transitions During Action Potentials Are Self-Similar to Each Other and to Variability of AP Firing Intervals Across the Broad Physiologic Range of AP Intervals During Autonomic Receptor Stimulation. Frontiers in Physiology, 2021. 12(953).

4. Jose, A.D. and D. Collison, The normal range and determinants of the intrinsic heart rate in man. Cardiovasc Res, 1970. 4(2): p. 160–7.

5. Freeling, J.L. and Y. Li, Age-related attenuation of parasympathetic control of the heart in mice. Int J Physiol Pathophysiol Pharmacol, 2015. 7(3): p. 126–35.

6. Whitehead, J.C., et al., A clinical frailty index in aging mice: comparisons with frailty index data in humans. J Gerontol A Biol Sci Med Sci, 2014. 69(6): p. 621–32.

7. Yaniv, Y., et al., deterioration of autonomic neuronal receptor signaling and mechanisms intrinsic to heart pacemaker cells contribute to age-associated alterations in heart rate variability in vivo. Aging Cell, 2016. 15(4): p. 716–24.

8. Dorey, T.W., et al., Impacts of frailty on heart rate variability in aging mice: Roles of the autonomic nervous system and sinoatrial node. Heart Rhythm, 2021.

9. Peters, C.H., E.J. Sharpe, and C. Proenza, Cardiac Pacemaker Activity and Aging. Annu Rev Physiol, 2020. 82: p. 21–43.

10. Moghtadaei, M., et al., The impacts of age and frailty on heart rate and sinoatrial node function. J Physiol, 2016. 594(23): p. 7105–7126.

11. Howlett, S.E. and K. Rockwood, Factors that influence reliability of the mouse clinical frailty index. J Gerontol A Biol Sci Med Sci, 2015. 70(6): p. 696.

12. Behar, J.A., et al., PhysioZoo: A Novel Open Access Platform for Heart Rate Variability Analysis of Mammalian Electrocardiographic Data. Front Physiol, 2018. 9: p. 1390.

13. Team, R.C., R: A language and environment for statistical computing. 2020.

14. RStudio Team, RStudio: Integrated Development for R. 2020, RStudio.

15. Wickham, H., ggplot2 : Elegant Graphics for Data Analysis, in Use R!,. 2016, Springer International Publishing : Imprint: Springer,: Cham. p. XVI, 260 pages 232 illustrations, 140 illustrations in color.

16. Kane, A.E., et al., Age, Sex and Overall Health, Measured As Frailty, Modify Myofilament Proteins in Hearts From Naturally Aging Mice. Sci Rep, 2020. 10(1): p. 10052.

17. Petr, M.A., et al., A cross-sectional study of functional and metabolic changes during aging through the lifespan in male mice. Elife, 2021. 10.

18. Costa MD, Davis RB, Goldberger AL. Heart Rate Fragmentation: A New Approach to the Analysis of Cardiac Interbeat Interval Dynamics. Frontiers in Physiology. 2017 May 9; 8:1–13. DOI=10.3389/fphys.2017.00255

19. Costa MD, Goldberger AL. Heart rate fragmentation: using cardiac pacemaker dynamics to probe the pace of biological aging. Am J Physiol Heart Circ Physiol. 2019 Jun 1;316(6):H1341–H1344.

20. Shaffer, F., R. McCraty, and C.L. Zerr, A healthy heart is not a metronome: an integrative review of the heart’s anatomy and heart rate variability. Front Psychol, 2014. 5: p. 1040.

21. Peng, C.K., et al., Quantification of scaling exponents and crossover phenomena in nonstationary heartbeat time series. Chaos, 1995. 5(1): p. 82–7.

22. Piskorski, J. and P. Guzik, Filtering Poincaré plots. computational methods in science and technology, 2005. 11: p. 39–48.

23. Hayano, J., et al., Accuracy of assessment of cardiac vagal tone by heart rate variability in normal subjects. Am J Cardiol, 1991. 67(2): p. 199–204.

24. Bhargava, V. and A.L. Goldberger, Effect of exercise in healthy men on QRS power spectrum. Am J Physiol, 1982. 243(6): p. H964–9.

25. Lensen, I.S., et al., Heart rate fragmentation gives novel insights into non-autonomic mechanisms governing beat-to-beat control of the heart’s rhythm. JRSM Cardiovasc Dis, 2020. 9: p. 2048004020948732.

26. Lakatta, E.G., Deficient neuroendocrine regulation of the cardiovascular system with advancing age in healthy humans. Circulation, 1993. 87(2): p. 631–6.

27. Lakatta, E.G., Cardiovascular regulatory mechanisms in advanced age. Physiol Rev, 1993. 73(2): p. 413–67.

28. Alley, D.E., et al., Changes in weight at the end of life: characterizing weight loss by time to death in a cohort study of older men. Am J Epidemiol, 2010. 172(5): p. 558–65.

29. Davies, L.C., et al., Enhanced prognostic value from cardiopulmonary exercise testing in chronic heart failure by non-linear analysis: oxygen uptake efficiency slope. Eur Heart J, 2006. 27(6): p. 684–90.

30. Laine, H., et al., Myocardial oxygen consumption is unchanged but efficiency is reduced in patients with essential hypertension and left ventricular hypertrophy. Circulation, 1999. 100(24): p. 2425–30.

31. Lakatta, E.G., Heartbeat music. Heart Rhythm, 2021. 18(5): p. 811–812.

32. Tagirova Sirenko, S., et al., Self-Similar Synchronization of Calcium and Membrane Potential Transitions During Action Potential Cycles Predict Heart Rate Across Species. JACC Clin Electrophysiol, 2021.

33. Liu, J., et al., Age-associated abnormalities of intrinsic automaticity of sinoatrial nodal cells are linked to deficient cAMP-PKA-Ca(2+) signaling. Am J Physiol Heart Circ Physiol, 2014. 306(10): p. H1385–97.

34. Weiss, J.N., et al., Early afterdepolarizations and cardiac arrhythmias. Heart Rhythm, 2010. 7(12): p. 1891–9.

35. Lakatta, E.G., V.A. Maltsev, and T.M. Vinogradova, A coupled SYSTEM of intracellular Ca2+ clocks and surface membrane voltage clocks controls the timekeeping mechanism of the heart’s pacemaker. Circ Res, 2010. 106(4): p. 659–73.

36. Moen, J.M., et al., Overexpression of a Neuronal Type Adenylyl Cyclase (Type 8) in Sinoatrial Node Markedly Impacts Heart Rate and Rhythm. Front Neurosci, 2019. 13: p. 615.

37. Lakatta, E.G., Beyond Bowditch: the convergence of cardiac chronotropy and inotropy. Cell Calcium, 2004. 35(6): p. 629–42.

38. Monfredi, O. and E.G. Lakatta, Complexities in cardiovascular rhythmicity: perspectives on circadian normality, ageing and disease. Cardiovasc Res, 2019. 115(11): p. 1576–1595.

39. Yaniv, Y., et al., Synchronization of sinoatrial node pacemaker cell clocks and its autonomic modulation impart complexity to heart beating intervals. Heart Rhythm, 2014. 11(7): p. 1210–9.

40. Costa, M.D., R.B. Davis, and A.L. Goldberger, Heart Rate Fragmentation: A Symbolic Dynamical Approach. Front Physiol, 2017. 8: p. 827.

41. Stein, P.K., et al., Development of more erratic heart rate patterns is associated with mortality postmyocardial infarction. J Electrocardiol, 2008. 41(2): p. 110–5.

42. Fried, L.P., et al., Untangling the concepts of disability, frailty, and comorbidity: implications for improved targeting and care. J Gerontol A Biol Sci Med Sci, 2004. 59(3): p. 255–63.

