## Supplement for "Emergence of Heartbeat Frailty in Advanced Age: Perspectives from Life-Long EKG Recordings in Mice"

Journal: GeroScience

Authors: Jack M Moen, Christopher H. Morrell, Ismayil Ahmet, Michael G. Matt, Moran Davoodi, Michael Petr, Shaquille Charles, Raphael deCabo, Yael Yaniv, and Edward G Lakatta

**Corresponding Author**:

Edward Lakatta, LCS, NIA, NIH

Supplement Table S1A. Descriptive statistics of HR and HRV parameters in Long Lived mice.

|  |  | Intrinsic State (during double autonomic blockade) | | | | | | | | |
| --- | --- | --- | --- | --- | --- | --- | --- | --- | --- | --- |
|  | Age | 6 | 9 | 12 | 15 | 18 | 21 | 24 | 27 | 30 |
| **Time Domain** | |  |  |  |  |  |  |  |  |  |
|  | n^*^ | 30 | 28 | 29 | 30 | 28 | 29 | 28 | 16 | 3 |
| Mean RR (ms) | mean | 135.86 | 133.81 | 142.93 | 131.73 | 139.33 | 138.09 | 151.32 | 196.1 | 154.75 |
|  | sd | 7.5543 | 6.9557 | 6.3442 | 6.7944 | 10.108 | 9.7206 | 9.8015 | 14.478 | 69.227 |
|  | n | 30 | 28 | 29 | 30 | 28 | 29 | 28 | 16 | 3 |
| SD RR (ms) | mean | 1.91 | 1.616 | 1.8154 | 1.5356 | 2.1781 | 1.802 | 2.2878 | 5.1416 | 29.159 |
|  | sd | 0.5472 | 0.5025 | 0.604 | 0.6912 | 2.306 | 0.8568 | 1.3248 | 3.585 | 40.273 |
| **Frequency Domain** | | |  |  |  |  |  |  |  |  |
| Total Power PSD  (ms^2^) | n | 30 | 28 | 29 | 30 | 28 | 29 | 28 | 16 | 3 |
|  | mean | 1.7269 | 1.2817 | 1.6601 | 1.3501 | 10.286 | 2.2217 | 5.43 | 19.707 | 44.669 |
|  | sd | 1.0202 | 0.7197 | 1.0052 | 1.6521 | 34.37 | 2.6958 | 12.486 | 23.287 | 37.742 |
|  | n | 30 | 28 | 29 | 30 | 28 | 29 | 28 | 16 | 3 |
| VLF (%) | mean | 8.5982 | 10.944 | 14.587 | 17.682 | 15.438 | 15.317 | 18.881 | 29.891 | 40.329 |
|  | sd | 4.6037 | 8.8516 | 13.031 | 12.701 | 10.15 | 10.48 | 10.178 | 15.225 | 30.462 |
|  | n | 30 | 28 | 29 | 30 | 28 | 29 | 28 | 16 | 3 |
| LF (%) | mean | 8.9183 | 10.891 | 12.756 | 11.72 | 14.401 | 16.543 | 13.281 | 21.088 | 65.082 |
|  | sd | 5.8251 | 6.2324 | 9.6027 | 7.3111 | 12.266 | 10.575 | 11.947 | 13.919 | 31.832 |
|  | n | 30 | 28 | 29 | 30 | 28 | 29 | 28 | 16 | 3 |
| HF (%) | mean | 77.513 | 72.832 | 65.299 | 62.606 | 59.341 | 53.457 | 48.169 | 26.461 | 47.794 |
|  | sd | 10.752 | 15.446 | 20.821 | 20.871 | 22.597 | 22.212 | 22.361 | 20.61 | 45.237 |
|  | n | 30 | 28 | 29 | 30 | 28 | 29 | 28 | 16 | 3 |
| LF_TO_HF Ratio | mean | 0.1327 | 0.1759 | 0.2616 | 0.2495 | 0.2704 | 0.3572 | 0.5631 | 2.1338 | 27.815 |
|  | sd | 0.1024 | 0.1239 | 0.2192 | 0.179 | 0.2061 | 0.2518 | 1.2613 | 4.2574 | 41.361 |
|  | n | 30 | 28 | 29 | 30 | 28 | 29 | 28 | 16 | 3 |
| β (N.U.) | mean | -1.383 | -1.315 | -1.233 | -1.371 | -1.325 | -1.231 | -1.289 | -1.282 | 24.405 |
|  | sd | 0.22 | 0.2014 | 0.3093 | 0.2907 | 0.3397 | 0.3892 | 0.3999 | 0.5426 | 44.214 |
| **Non-linear Domain** | | |  |  |  |  |  |  |  |  |
|  | n | 30 | 28 | 29 | 30 | 28 | 29 | 28 | 16 | 3 |
| SD1 (ms) | mean | 2.0242 | 1.7053 | 1.8506 | 1.4973 | 2.1036 | 1.6488 | 2.0917 | 3.7887 | 27.904 |
|  | sd | 0.6206 | 0.5874 | 0.7486 | 0.8172 | 2.4967 | 0.8243 | 1.4499 | 3.3762 | 41.252 |
|  | n | 30 | 28 | 29 | 30 | 28 | 29 | 28 | 16 | 3 |
| SD2 (ms) | mean | 1.769 | 1.5027 | 1.7363 | 1.5254 | 2.0875 | 1.861 | 2.3579 | 5.7731 | 30.013 |
|  | sd | 0.4923 | 0.4351 | 0.5198 | 0.6275 | 1.9782 | 0.9276 | 1.2871 | 3.9549 | 39.644 |
|  | n | 30 | 28 | 29 | 30 | 28 | 29 | 28 | 16 | 3 |
| SD Ratio | mean | 1.1393 | 1.114 | 1.0434 | 0.9656 | 0.9819 | 0.8964 | 0.8476 | 0.6729 | 0.7431 |
|  | sd | 0.1411 | 0.2005 | 0.2751 | 0.2686 | 0.2441 | 0.2751 | 0.2726 | 0.3452 | 0.2346 |
|  | n | 30 | 28 | 29 | 30 | 28 | 29 | 28 | 16 | 3 |
| α_1_ (N.U.) | mean | 0.267 | 0.3123 | 0.3605 | 0.3713 | 0.3633 | 0.4176 | 0.3727 | 0.531 | 25.642 |
|  | sd | 0.1011 | 0.1207 | 0.166 | 0.1412 | 0.1132 | 0.1439 | 0.1956 | 0.3097 | 43.143 |
|  | n | 30 | 28 | 29 | 30 | 28 | 29 | 28 | 16 | 3 |
| α_2_ (N.U.) | mean | 0.5837 | 0.6022 | 0.6663 | 0.6865 | 0.6718 | 0.6429 | 0.7434 | 0.8097 | 25.578 |
|  | sd | 0.1465 | 0.1759 | 0.203 | 0.2273 | 0.1965 | 0.1916 | 0.1883 | 0.2572 | 43.198 |
| Sample Entropy  (N.U.) | n | 30 | 28 | 29 | 30 | 28 | 29 | 28 | 16 | 3 |
|  | mean | 1.3289 | 1.3638 | 1.3767 | 1.4454 | 1.447 | 1.4457 | 1.332 | 1.0989 | 25.79 |
|  | sd | 0.2012 | 0.2971 | 0.2781 | 0.3429 | 0.3453 | 0.2827 | 0.3164 | 0.4287 | 43.016 |

| **Heart Rate Fragmentation** | | | |  |  |  |  |  |  |  |
| --- | --- | --- | --- | --- | --- | --- | --- | --- | --- | --- |
|  | n | 30 | 28 | 29 | 30 | 28 | 29 | 28 | 16 | 3 |
| PIP (%) | mean | 56.331 | 59.513 | 61.148 | 59.923 | 60.791 | 59.672 | 59.179 | 59.168 | 60.231 |
|  | sd | 4.8508 | 5.1295 | 4.6558 | 4.3934 | 4.9786 | 4.3829 | 5.2608 | 5.1327 | 17.504 |
|  | n | 30 | 28 | 29 | 30 | 28 | 29 | 28 | 16 | 3 |
| IALS (N.U.) | mean | 0.5645 | 0.5963 | 0.6128 | 0.6004 | 0.6092 | 0.598 | 0.5931 | 0.5935 | 25.505 |
|  | sd | 0.0486 | 0.0513 | 0.0466 | 0.044 | 0.0499 | 0.0439 | 0.0527 | 0.0514 | 43.261 |
|  | n | 30 | 28 | 29 | 30 | 28 | 29 | 28 | 16 | 3 |
| PSS (%) | mean | 66.12 | 73.606 | 75.775 | 74.967 | 74.546 | 69.684 | 62.799 | 63.905 | 60.173 |
|  | sd | 11.908 | 13.117 | 10.695 | 9.0841 | 10.007 | 11.084 | 15.702 | 7.9964 | 21.741 |
|  | n | 30 | 28 | 29 | 30 | 28 | 29 | 28 | 16 | 3 |
| PAS (%) | mean | 10.326 | 12.559 | 13.706 | 11.966 | 14.555 | 14.479 | 20.067 | 27.342 | 40.851 |
|  | sd | 7.1928 | 10.623 | 11.989 | 9.0871 | 11.381 | 9.5201 | 10.646 | 7.7683 | 31.79 |

Supplement Table S1B. Descriptive statistics of HR and HRV parameters in Long Lived mice.

|  |  | Basal State | | | | | | | | |
| --- | --- | --- | --- | --- | --- | --- | --- | --- | --- | --- |
|  | Age | 6 | 9 | 12 | 15 | 18 | 21 | 24 | 27 | 30 |
| **Time Domain** |  |  |  |  |  |  |  |  |  |  |
|  | n^*^ | 30 | 28 | 29 | 30 | 28 | 29 | 28 | 17 | 3 |
| Mean RR (ms) | mean | 135.8 | 135.7 | 140.8 | 136.3 | 136.2 | 138.4 | 139.6 | 151.2 | 147.3 |
|  | sd | 8.234 | 9.498 | 10.56 | 11.23 | 13.63 | 12.74 | 14.19 | 15.59 | 22.16 |
|  | n | 30 | 28 | 29 | 30 | 28 | 29 | 28 | 17 | 3 |
| SD RR (ms) | mean | 2.876 | 3.146 | 2.895 | 2.816 | 2.83 | 3.159 | 3.697 | 4.504 | 4.826 |
|  | sd | 1.128 | 2.352 | 1.186 | 1.523 | 1.777 | 2.009 | 2.86 | 3.176 | 5.607 |
| **Frequency Domain** |  |  |  |  |  |  |  |  |  |  |
|  | n | 30 | 28 | 29 | 30 | 28 | 29 | 28 | 17 | 3 |
| Total Power PSD | mean | 4.224 | 6.77 | 4.571 | 4.578 | 5.15 | 6.646 | 11.09 | 15.46 | 29.69 |
| (ms^2^) | sd | 3.053 | 14.23 | 3.708 | 5.816 | 9.062 | 9.796 | 20.68 | 19.38 | 48.39 |
|  | n | 30 | 28 | 29 | 30 | 28 | 29 | 28 | 17 | 3 |
| VLF (%) | mean | 28.06 | 33.73 | 33.41 | 35.48 | 36.04 | 33.74 | 36.29 | 26.1 | 25.86 |
|  | sd | 11.38 | 13.6 | 14.43 | 14.2 | 17.82 | 16.9 | 17.74 | 15.79 | 1.46 |
|  | n | 30 | 28 | 29 | 30 | 28 | 29 | 28 | 17 | 3 |
| LF (%) | mean | 8.607 | 7.895 | 10.63 | 7.737 | 8.464 | 13.53 | 11.7 | 26.54 | 51.58 |
|  | sd | 7.211 | 4.228 | 9.871 | 5.082 | 7.682 | 12.78 | 10.15 | 15.2 | 18.1 |
|  | n | 30 | 28 | 29 | 30 | 28 | 29 | 28 | 17 | 3 |
| HF (%) | mean | 47.77 | 41.38 | 41.24 | 35.73 | 37.9 | 34.9 | 34.32 | 42.16 | 18.32 |
|  | sd | 16.12 | 17.7 | 22.16 | 20.53 | 21.96 | 19.66 | 23.73 | 27.57 | 8.872 |
|  | n | 30 | 28 | 29 | 30 | 28 | 29 | 28 | 17 | 3 |
| LF_TO_HF Ratio | mean | 0.207 | 0.329 | 0.427 | 0.561 | 0.345 | 0.502 | 0.509 | 1.41 | 3.731 |
|  | sd | 0.22 | 0.525 | 0.654 | 1.483 | 0.366 | 0.543 | 0.534 | 1.651 | 3.04 |
|  | n | 30 | 28 | 29 | 30 | 28 | 29 | 28 | 17 | 3 |
| β (N.U.) | mean | -1.64 | -1.66 | -1.5 | -1.71 | -1.67 | -1.46 | -1.35 | -1.06 | -0.97 |
|  | sd | 0.197 | 0.234 | 0.353 | 0.168 | 0.251 | 0.504 | 0.529 | 0.463 | 0.722 |
| **Non-linear Domain** |  |  |  |  |  |  |  |  |  |  |
|  | n | 30 | 28 | 29 | 30 | 28 | 29 | 28 | 17 | 3 |
| SD1 (ms) | mean | 2.261 | 2.18 | 2.218 | 2.014 | 2.069 | 2.063 | 2.738 | 3.591 | 2.493 |
|  | sd | 0.964 | 1.148 | 1.204 | 1.56 | 1.694 | 1.176 | 3.013 | 3.033 | 2.646 |
|  | n | 30 | 28 | 29 | 30 | 28 | 29 | 28 | 17 | 3 |
| SD2 (ms) | mean | 3.317 | 3.781 | 3.324 | 3.309 | 3.32 | 3.827 | 4.238 | 4.892 | 6.338 |
|  | sd | 1.364 | 3.225 | 1.372 | 1.709 | 2.024 | 2.664 | 2.959 | 3.723 | 7.468 |
|  | n | 30 | 28 | 29 | 30 | 28 | 29 | 28 | 17 | 3 |
| SD Ratio | mean | 0.703 | 0.649 | 0.68 | 0.596 | 0.632 | 0.591 | 0.597 | 0.81 | 0.442 |
|  | sd | 0.195 | 0.229 | 0.291 | 0.265 | 0.276 | 0.254 | 0.331 | 0.412 | 0.064 |
|  | n | 30 | 28 | 29 | 30 | 28 | 29 | 28 | 17 | 3 |
| α_1_ (N.U.) | mean | 0.346 | 0.394 | 0.443 | 0.427 | 0.4 | 0.501 | 0.496 | 0.599 | 0.975 |
|  | sd | 0.147 | 0.224 | 0.235 | 0.245 | 0.205 | 0.206 | 0.247 | 0.3 | 0.285 |
|  | n | 30 | 28 | 29 | 30 | 28 | 29 | 28 | 17 | 3 |
| α_2_ (N.U.) | mean | 1.008 | 1.097 | 1.041 | 1.096 | 1.088 | 1.035 | 1.081 | 0.742 | 0.616 |
|  | sd | 0.203 | 0.13 | 0.2 | 0.216 | 0.257 | 0.268 | 0.271 | 0.255 | 0.155 |
|  | n | 30 | 28 | 29 | 30 | 28 | 29 | 28 | 17 | 3 |
| Sample Entropy | mean | 1.468 | 1.449 | 1.436 | 1.389 | 1.412 | 1.402 | 1.34 | 1.328 | 1.353 |
| (N.U.) | sd | 0.301 | 0.372 | 0.287 | 0.33 | 0.31 | 0.319 | 0.339 | 0.399 | 0.511 |

| **Heart Rate Fragmentation** | |  |  |  |  |  |  |  |  |  |
| --- | --- | --- | --- | --- | --- | --- | --- | --- | --- | --- |
|  | n | 30 | 28 | 29 | 30 | 28 | 29 | 28 | 17 | 3 |
| PIP (%) | mean | 56.14 | 58.85 | 60.29 | 61.08 | 60.49 | 60.08 | 56.8 | 56.9 | 45.24 |
|  | sd | 4.899 | 4.479 | 3.975 | 4.816 | 5.397 | 4.215 | 5.473 | 7.813 | 3.703 |
|  | n | 30 | 28 | 29 | 30 | 28 | 29 | 28 | 17 | 3 |
| IALS (N.U.) | mean | 0.563 | 0.59 | 0.604 | 0.612 | 0.606 | 0.602 | 0.569 | 0.57 | 0.454 |
|  | sd | 0.049 | 0.045 | 0.04 | 0.048 | 0.054 | 0.042 | 0.055 | 0.078 | 0.037 |
|  | n | 30 | 28 | 29 | 30 | 28 | 29 | 28 | 17 | 3 |
| PSS (%) | mean | 74.53 | 79.53 | 81.49 | 79.9 | 78.64 | 76.43 | 67.73 | 61.76 | 42.35 |
|  | sd | 9.979 | 10.26 | 10.3 | 12.43 | 11.7 | 10.68 | 12.42 | 12.81 | 5.667 |
|  | n | 30 | 28 | 29 | 30 | 28 | 29 | 28 | 17 | 3 |
| PAS (%) | mean | 11.16 | 10.86 | 10.29 | 14.83 | 14.18 | 12.95 | 15.9 | 23.88 | 10.42 |
|  | sd | 9.803 | 9.033 | 8.032 | 10.37 | 13.53 | 7.534 | 11.49 | 12.48 | 5.318 |

Supplement Table S1C. Descriptive statistics of HR and HRV parameters in Long Lived mice.

|  | Basal - Intrinsic = Effect of Autonomic Input | | | | | | | | |
| --- | --- | --- | --- | --- | --- | --- | --- | --- | --- |
| Age | 6 | 9 | 12 | 15 | 18 | 21 | 24 | 27 | 30 |
| **Time Domain** |  |  |  |  |  |  |  |  |  |
|  | 30 | 28 | 29 | 30 | 28 | 29 | 28 | 16 | 3 |
| Mean RR (ms) | -0.04 | 1.85 | -2.18 | 4.57 | -3.13 | 0.27 | -11.70 | -46.52 | -7.42 |
|  | 5.72 | 8.34 | 7.80 | 8.57 | 11.40 | 7.68 | 8.26 | 8.10 | 85.35 |
|  | 30 | 28 | 29 | 30 | 28 | 29 | 28 | 16 | 3 |
| SD RR (ms) | 0.97 | 1.53 | 1.08 | 1.28 | 0.65 | 1.36 | 1.41 | -0.72 | -24.33 |
|  | 1.04 | 2.18 | 0.94 | 1.09 | 1.64 | 1.58 | 2.22 | 4.39 | 34.89 |
| **Frequency Domain** | |  |  |  |  |  |  |  |  |
|  | 30 | 28 | 29 | 30 | 28 | 29 | 28 | 16 | 3 |
| Total Power PSD | 2.50 | 5.49 | 2.91 | 3.23 | -5.14 | 4.42 | 5.66 | -4.91 | -14.98 |
| (ms^2^) | 2.84 | 14.07 | 3.27 | 4.58 | 33.22 | 8.54 | 10.83 | 30.19 | 35.61 |
|  | 30 | 28 | 29 | 30 | 28 | 29 | 28 | 16 | 3 |
| VLF (%) | 19.46 | 22.78 | 18.82 | 17.80 | 20.60 | 18.43 | 17.41 | -4.44 | -14.47 |
|  | 12.53 | 13.50 | 11.68 | 13.69 | 16.01 | 14.56 | 15.47 | 16.63 | 31.76 |
|  | 30 | 28 | 29 | 30 | 28 | 29 | 28 | 16 | 3 |
| LF (%) | -0.31 | -3.00 | -2.12 | -3.98 | -5.94 | -3.01 | -1.58 | 6.21 | -13.50 |
|  | 8.83 | 5.53 | 8.89 | 6.16 | 10.76 | 8.84 | 8.14 | 22.54 | 25.71 |
|  | 30 | 28 | 29 | 30 | 28 | 29 | 28 | 16 | 3 |
| HF (%) | -29.74 | -31.46 | -24.06 | -26.88 | -21.44 | -18.56 | -13.85 | 18.07 | -29.47 |
|  | 20.64 | 17.23 | 20.16 | 22.50 | 26.98 | 26.14 | 28.10 | 22.61 | 53.10 |
|  | 30 | 28 | 29 | 30 | 28 | 29 | 28 | 16 | 3 |
| LF_TO_HF Ratio | 0.07 | 0.15 | 0.17 | 0.31 | 0.07 | 0.14 | -0.05 | -1.05 | -24.08 |
|  | 0.24 | 0.49 | 0.59 | 1.38 | 0.34 | 0.47 | 1.16 | 4.56 | 38.54 |
|  | 30 | 28 | 29 | 30 | 28 | 29 | 28 | 16 | 3 |
| β (N.U.) | -0.261 | -0.341 | -0.268 | -0.336 | -0.348 | -0.228 | -0.058 | 0.228 | -25.378 |
|  | 0.294 | 0.307 | 0.370 | 0.303 | 0.346 | 0.392 | 0.550 | 0.687 | 43.528 |
| **Non-linear Domain** | |  |  |  |  |  |  |  |  |
|  | 30 | 28 | 29 | 30 | 28 | 29 | 28 | 16 | 3 |
| SD1 (ms) | 0.236 | 0.474 | 0.367 | 0.517 | -0.034 | 0.414 | 0.646 | -0.104 | -25.411 |
|  | 0.884 | 0.859 | 0.848 | 0.955 | 1.352 | 0.854 | 2.627 | 3.173 | 38.714 |
|  | 30 | 28 | 29 | 30 | 28 | 29 | 28 | 16 | 3 |
| SD2 (ms) | 1.548 | 2.278 | 1.588 | 1.784 | 1.233 | 1.966 | 1.880 | -1.073 | -23.675 |
|  | 1.284 | 3.085 | 1.182 | 1.365 | 1.792 | 2.181 | 2.172 | 5.404 | 32.533 |
|  | 30 | 28 | 29 | 30 | 28 | 29 | 28 | 16 | 3 |
| SD Ratio | -0.437 | -0.465 | -0.363 | -0.370 | -0.350 | -0.306 | -0.251 | 0.171 | -0.301 |
|  | 0.239 | 0.209 | 0.249 | 0.256 | 0.306 | 0.311 | 0.335 | 0.357 | 0.295 |
|  | 30 | 28 | 29 | 30 | 28 | 29 | 28 | 16 | 3 |
| α_1_ (N.U.) | 0.079 | 0.082 | 0.082 | 0.056 | 0.037 | 0.084 | 0.123 | 0.031 | -24.667 |
|  | 0.176 | 0.208 | 0.182 | 0.191 | 0.181 | 0.206 | 0.217 | 0.401 | 42.873 |
|  | 30 | 28 | 29 | 30 | 28 | 29 | 28 | 16 | 3 |
| α_2_ (N.U.) | 0.424 | 0.495 | 0.374 | 0.410 | 0.416 | 0.392 | 0.337 | -0.062 | -24.962 |
|  | 0.226 | 0.203 | 0.169 | 0.282 | 0.290 | 0.238 | 0.262 | 0.248 | 43.302 |
|  | 30 | 28 | 29 | 30 | 28 | 29 | 28 | 16 | 3 |
| Sample Entropy | 0.14 | 0.09 | 0.06 | -0.06 | -0.03 | -0.04 | -0.01 | 0.26 | -24.44 |
| (N.U.) | 0.23 | 0.36 | 0.30 | 0.38 | 0.36 | 0.32 | 0.34 | 0.60 | 43.53 |

| **Heart Rate Fragmentation** | | |  |  |  |  |  |  |  |
| --- | --- | --- | --- | --- | --- | --- | --- | --- | --- |
|  | 30 | 28 | 29 | 30 | 28 | 29 | 28 | 16 | 3 |
| PIP (%) | -0.19 | -0.66 | -0.86 | 1.15 | -0.30 | 0.41 | -2.38 | -1.42 | -14.99 |
|  | 4.35 | 4.66 | 4.98 | 4.59 | 4.85 | 4.71 | 5.57 | 9.61 | 20.88 |
|  | 30 | 28 | 29 | 30 | 28 | 29 | 28 | 16 | 3 |
| IALS (N.U.) | -0.002 | -0.007 | -0.009 | 0.012 | -0.003 | 0.004 | -0.024 | -0.015 | -25.051 |
|  | 0.044 | 0.047 | 0.050 | 0.046 | 0.049 | 0.047 | 0.056 | 0.096 | 43.275 |
|  | 30 | 28 | 29 | 30 | 28 | 29 | 28 | 16 | 3 |
| PSS (%) | 8.41 | 5.92 | 5.71 | 4.93 | 4.10 | 6.75 | 4.93 | -0.72 | -17.82 |
|  | 13.55 | 11.16 | 9.88 | 7.60 | 10.75 | 10.72 | 13.61 | 16.80 | 26.93 |
|  | 30 | 28 | 29 | 30 | 28 | 29 | 28 | 16 | 3 |
| PAS (%) | 0.84 | -1.70 | -3.41 | 2.87 | -0.37 | -1.53 | -4.17 | -2.83 | -30.43 |
|  | 9.14 | 11.92 | 11.05 | 10.11 | 10.63 | 10.13 | 10.07 | 14.11 | 33.47 |

**Supplement Table S2. Statistical Analyses (*) the effects of age on Basal, Intrinsic, and Autonomic Input on Intrinsic RR interval mean and RR interval variability in long lived mice.**

|  | Intrinsic State | | Basal State | | Effect of Autonomic  Input on Intrinsic | |
| --- | --- | --- | --- | --- | --- | --- |
| Age (months) | 6 to 21 | 21 to 30 | 6 to 21 | 21 to 30 | 6 to 21 | 21 to 30 |
| Variable |  |  |  |  |  |  |
| Body Weight (g) |  |  | < 2.2e-16 ↑ | 3.5e-15 ↓ |  |  |
| **Time Domain** |  |  |  |  |  |  |
| Mean RR (ms) | 1.51E-06 ∽ | < 2.2e-16 ↑ |  | 0.01172 ↑ | 0.0008 ∽ | < 2.2e-16 ↑ |
| SD RR (ms) |  | 3.22e-6 ↑ |  |  |  | 0.0748 ↑ |
| **Frequency Domain** |  |  |  |  |  |  |
| Total Power PSD (ms^2^) |  | 0.0005 ↑ |  | 0.0984 ↑ |  |  |
| VLF (%) | 0.0008 ↑ | 0.00577 ↑ |  |  |  | 1.31e-5 ↑ |
| LF (%) | 0.0021 ↑ | 0.0034 ↑ | 0.0385 ↑ | 1.04e-6 ↑ |  | 0.0576 ↓↑ |
| HF (%) | 7.33e-7 ↓ | 4.35e-5 ↓ | 0.0657 ↓↑ |  |  | 4.35e-05 ↓ |
| LF TO HF Ratio | 9.00e-6 ↑ | 0.0047 ↑ |  | 3.97e-6 ↑ |  |  |
| β (N.U.) |  |  | 0.0071 ↑↓ | 0.05417 ↑ |  | 0.0316 ↓ |
| **Non-linear Domain** |  |  |  |  |  |  |
| SD1 (ms) |  | 0.0044 ↑ |  |  |  |  |
| SD2 (ms) |  | 5.39e-7 ↑ |  |  |  | 0.0317 ↑ |
| SDRatio | 3.95e-5 ↓ | 0.0253 ↓ |  | 0.0772 ∽ |  | 7.31e-5 ↓ |
| α_1_ (N.U.) | 1.77e-5 ↑ | 0.0059 ↑ | 0.0355 ↑ | 0.0120 ↑ |  |  |
| α_2_ (N.U.) |  | 0.05697 ↑ |  | 7.29e-5 ↓ |  | 1.90e-7 ↑ |
| Sample Entropy (N.U.) |  | 0.00681 ↓ |  |  |  | 0.0793 ↓↑ |
| **Heart Rate Fragmentation** |  |  |  |  |  |  |
| PIP (%) | 0.0002 ↑→ |  | 6.15e-5 ↑ | 0.0002 ↓ |  | 0.0444 ↑ |
| IALS (N.U.) | 0.0002 ↑→ |  | 6.22e-5 ↑ | 0.0002 ↓ |  | 0.0429↑ |
| PSS (%) | 0.0007 ↑↓ | 0.0318 ↓ | 0.0336 ↑↓ | 3.61e-6 ↓ |  | 0.0284 ↑ |
| PAS (%) |  | 0.00037 ↑ |  | 0.0053 ↑ |  |  |

Notes:

* 1. p-values from repeated-measures mixed ANOVA models; 2. Effect of Autonomic Input on Intrinsic = Basal - Intrinsic in each mouse;

3. N.U. = no units. 4. ↑ variable is significantly increasing with age, ↓ variable is significantly decreasing with age, and ∽ the variable is significantly oscillating.

Table S3. Correlations and p-values of Mean RR with additional variables excluded from those that clustered with the increased mean RR interval in Fig. 6.

|  | Mean RR | SDRR | Total Power | LF | LF/HF | β | SD1 | SD2 | α_1_ | Sample Entropy | PIP | IALS | PSS |
| --- | --- | --- | --- | --- | --- | --- | --- | --- | --- | --- | --- | --- | --- |
| Mean RR : r |  | 0.2485 | 0.0963 | 0.0926 | 0.0893 | -0.0534 | 0.2391 | 0.2430 | -0.0258 | 0.1376 | 0.3537 | 0.3648 | 0.2809 |
| Mean RR : p |  | 0.1854 | 0.6129 | 0.6266 | 0.6390 | 0.7794 | 0.2031 | 0.1958 | 0.8922 | 0.4684 | 0.0552 | 0.0475 | 0.1327 |
| SDRR : r | 0.2587 |  | 0.5317 | 0.0957 | -0.0175 | 0.3991 | 0.9566 | 0.9692 | -0.1033 | -0.4317 | 0.0001 | 0.1899 | 0.1279 |
| SDRR : p | 0.1675 |  | 0.0025 | 0.6149 | 0.9269 | 0.0289 | 0.0000 | 0.0000 | 0.5869 | 0.0172 | 0.9997 | 0.3148 | 0.5006 |
| Total Power : r | 0.1554 | 0.9314 |  | 0.4895 | 0.3407 | 0.2988 | 0.4144 | 0.4752 | 0.2520 | -0.1372 | 0.0347 | 0.0505 | 0.0091 |
| Total Power : p | 0.4122 | 8.2E-14 |  | 0.0060 | 0.0654 | 0.1087 | 0.0228 | 0.0080 | 0.1792 | 0.4698 | 0.8556 | 0.7911 | 0.9619 |
| LF : r | -0.1159 | 0.3861 | 0.5291 |  | 0.8305 | 0.5644 | -0.0094 | 0.1119 | 0.7343 | -0.1685 | -0.0540 | -0.0202 | -0.1941 |
| LF : p | 0.5421 | 0.0351 | 0.0026 |  | 1.4E-08 | 0.0012 | 0.9608 | 0.5559 | 3.9E-06 | 0.3734 | 0.7767 | 0.9154 | 0.3041 |
| LF/HF : r | 0.0393 | 0.4427 | 0.6258 | 0.7216 |  | 0.3529 | -0.2171 | 0.1055 | 0.9259 | -0.1505 | 0.0260 | 0.0954 | -0.1583 |
| LF/HF : p | 0.8367 | 0.0143 | 0.0002 | 6.8E-06 |  | 0.0558 | 0.2492 | 0.5789 | 2.3E-13 | 0.4272 | 0.8917 | 0.6162 | 0.4035 |
| β : r | 0.0106 | 0.5640 | 0.6156 | 0.6493 | 0.5574 |  | 0.3978 | 0.3573 | 0.3180 | -0.4953 | -0.0080 | 0.1423 | 0.0889 |
| β : p | 0.9558 | 0.0012 | 0.0003 | 0.0001 | 0.0014 |  | 0.0295 | 0.0526 | 0.0868 | 0.0054 | 0.9667 | 0.4533 | 0.6404 |
| SD1 : r | 0.1446 | 0.8441 | 0.6910 | 0.2142 | 0.0355 | 0.4208 |  | 0.8685 | -0.2847 | -0.3777 | 0.0165 | 0.1527 | 0.1578 |
| SD1 : p | 0.4458 | 4.6E-09 | 2.4E-05 | 0.2558 | 0.8521 | 0.0206 |  | 5.0E-10 | 0.1273 | 0.0396 | 0.9310 | 0.4204 | 0.4051 |
| SD2 : r | 0.2798 | 0.9502 | 0.9531 | 0.4783 | 0.6192 | 0.5750 | 0.6451 |  | 0.0171 | -0.4415 | -0.0331 | 0.2089 | 0.0913 |
| SD2 : p | 0.1343 | 8.9E-16 | 4.4E-16 | 0.0075 | 0.0003 | 0.0009 | 0.0001 |  | 0.9285 | 0.0146 | 0.8620 | 0.2680 | 0.6313 |
| α_1_ : r | 0.0177 | 0.4036 | 0.5433 | 0.5485 | 0.8007 | 0.4574 | -0.0352 | 0.5920 |  | -0.1303 | 0.0024 | 0.0481 | -0.0751 |
| α_1_ : p | 0.9260 | 0.0270 | 0.0019 | 0.0017 | 1.1E-07 | 0.0110 | 0.8537 | 0.0006 |  | 0.4927 | 0.9899 | 0.8006 | 0.6931 |
| Sample Entropy : r | 0.1937 | -0.6270 | -0.5650 | -0.3840 | -0.3243 | -0.6726 | -0.5670 | -0.5778 | -0.3180 |  | -0.0095 | -0.1335 | -0.2226 |
| Sample Entropy : p | 0.3050 | 0.0002 | 0.0011 | 0.0362 | 0.0804 | 4.7E-05 | 0.0011 | 0.0008 | 0.0868 |  | 0.9604 | 0.4818 | 0.2371 |
| PIP : r | 0.1205 | -0.2876 | -0.3709 | -0.5315 | -0.4529 | -0.3071 | -0.1704 | -0.3343 | -0.3228 | 0.3053 |  | 0.8606 | 0.7326 |
| PIP : p | 0.5258 | 0.1233 | 0.0436 | 0.0025 | 0.0120 | 0.0988 | 0.3679 | 0.0710 | 0.0819 | 0.1009 |  | 1.1E-09 | 4.2E-06 |
| IALS : r | 0.1227 | -0.2867 | -0.3701 | -0.5316 | -0.4529 | -0.3070 | -0.1697 | -0.3334 | -0.3229 | 0.3055 | 1.0000 |  | 0.7631 |
| IALS : p | 0.5184 | 0.1245 | 0.0441 | 0.0025 | 0.0120 | 0.0989 | 0.3701 | 0.0718 | 0.0818 | 0.1006 | 0.0000 |  | 9.4E-07 |
| PSS : r | -0.1424 | 0.3475 | 0.3579 | 0.3711 | 0.4174 | 0.3070 | 0.3041 | 0.3144 | 0.2964 | -0.2977 | -0.6038 | -0.6043 |  |
| PSS : p | 0.4530 | 0.0599 | 0.0521 | 0.0435 | 0.0217 | 0.0989 | 0.1023 | 0.0907 | 0.1117 | 0.1101 | 0.0004 | 0.0004 |  |

Upper triangle: contains the correlations in the early time period (before 21 months); lower triangle contains the correlations in the late time period (after 21 months).

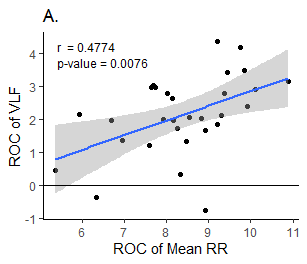

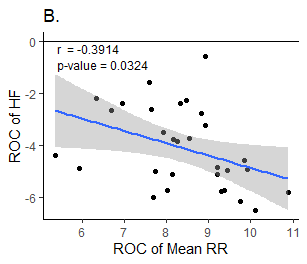

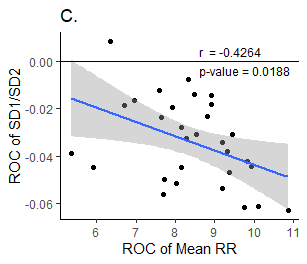

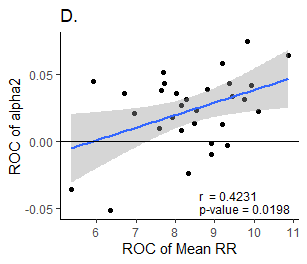

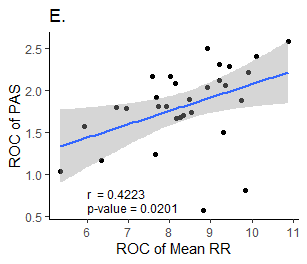

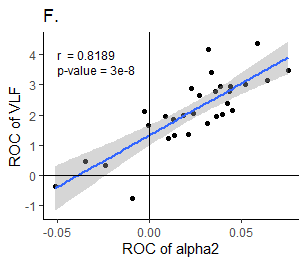

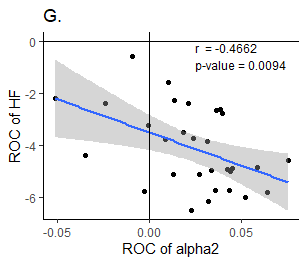

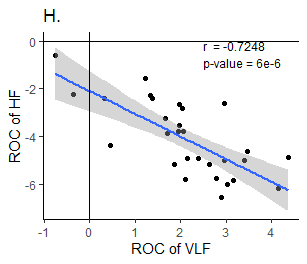

Figure S1. Associations among Intrinsic Rates of Change in long-lived mice.

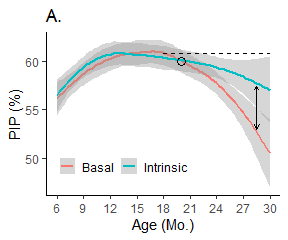

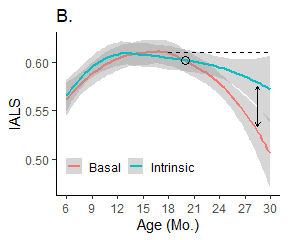

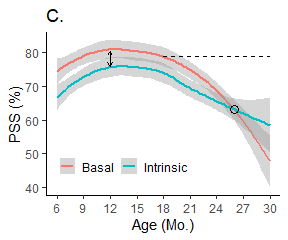

Figure S2. Loess curves for Heart Rate Fragmentation variables.

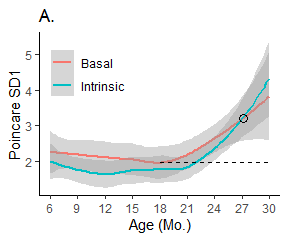

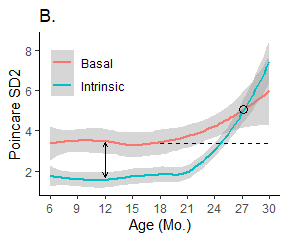

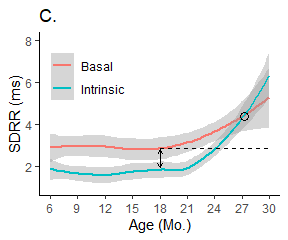

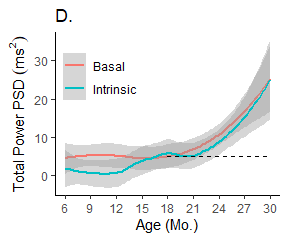

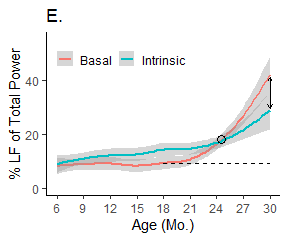

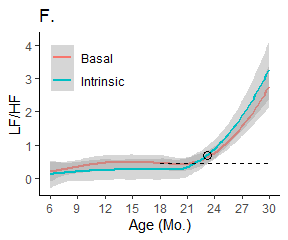

Supplement Figure S3. Average loess smooth curves for basal and intrinsic RR interval variablilty signatures and on the mean RR interval for variables excluded from the cluster abalysis during the enitre life course of all long-lived mice. Arrows show the effect of autonomic modulation.

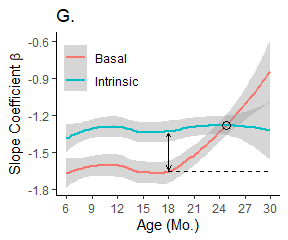

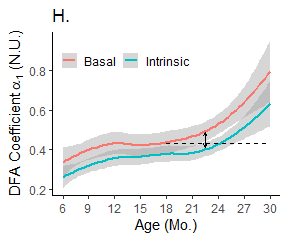

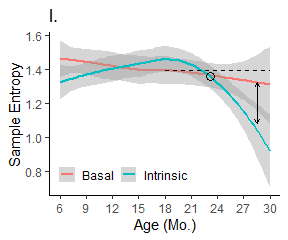

Supplement Figure S3 (continued).
